# Young transposable elements rewired gene regulatory networks in human and chimpanzee hippocampal intermediate progenitors

**DOI:** 10.1101/2021.11.24.469877

**Authors:** Sruti Patoori, Samantha M. Barnada, Christopher Large, John I. Murray, Marco Trizzino

## Abstract

The hippocampus is associated with essential brain functions such as learning and memory. Human hippocampal volume is significantly greater than expected when compared to non-human apes, suggesting a recent expansion. Intermediate progenitors, which are able to undergo multiple rounds of proliferative division before a final neurogenic division, may have played a role in the evolutionary hippocampal expansion. To investigate the evolution of gene regulatory networks underpinning hippocampal neurogenesis in apes, we leveraged the differentiation of human and chimpanzee induced Pluripotent Stem Cells into TBR2-positive hippocampal intermediate progenitors (hpIPCs). We find that the gene networks active in hpIPCs are significantly different between humans and chimpanzees, with ∼2,500 genes differentially expressed. We demonstrate that species-specific transposon-derived enhancers contribute to these transcriptomic differences. Young transposons, predominantly Endogenous Retroviruses (ERVs) and SINE-Vntr-Alus (SVAs), were co-opted as enhancers in a species-specific manner. Human-specific SVAs provided substrates for thousands of novel TBR2 binding sites, and CRISPR-mediated repression of these SVAs attenuates the expression of ∼25% of the genes that are upregulated in human intermediate progenitors relative to the same cell population in the chimpanzee.

**Summary statement:** Evolution of human and chimpanzee hippocampal development was mediated by co-option of young retrotransposons into species-specific enhancers.

## Introduction

The hippocampus is associated with many traits relevant in the context of human evolution. These include traits such as tool use and language which require social cognition and learning, as well as spatial memory, navigation, and episodic memory (Burgess et al., 2002; Eichenbaum, 2017a; Eichenbaum, 2017b; Squire, 1992; Tomasello and Herrmann, 2010). This region of the brain is also greatly affected by Alzheimer’s Disease (AD), a neurodegenerative disorder characterized by cell death, plaques and tangles of misfolded proteins, and cognitive decline (Duyckaerts et al., 2009). It has been hypothesized that the cognitive AD phenotype is uniquely human and that non-human primates, including chimpanzees, do not exhibit AD-related dementia (Edler et al., 2017; Finch and Austad, 2015; Walker and Jucker, 2017). If humans are uniquely susceptible to AD, it is crucial to understand how the human hippocampus differs from that of our closest biological relatives, the chimpanzees.

Human hippocampal volume is 50% greater than expected when compared to the hippocampal volumes of non-human apes, possibly indicating a recent hippocampal expansion specific to the human lineage (Barger et al., 2014). However, the evolution of the human hippocampus and the developmental mechanisms driving the human-specific volume increase have not yet been thoroughly studied.

Recent studies have suggested that evolutionary changes to neuronal progenitors may have had an impact on cortical volume in primates by increasing proliferative potential (Martínez-Cerdeño et al., 2006; Rétaux et al., 2013; Florio and Huttner, 2014). A specific class of neuronal progenitors known as intermediate progenitor cells (IPCs) or “transit amplifying cells” are able to undergo multiple rounds of proliferative division before a final neurogenic division (Englund, 2005; Arnold et al., 2008; Pontious et al., 2008; Hevner, 2019). These cells express the neurodevelopmental transcription factor TBR2 (EOMES) and are found in the sub-ventricular zone of the developing neocortex and hippocampus (Bulfone et al., 1999; Kimura et al., 1999; Englund, 2005; Cipriani et al., 2016). Genetic ablation of TBR2 in these progenitors results in reduced cortical thickness in mice (Sessa et al., 2008), abnormal cortical cell differentiation (Mihalas et al., 2016), and impaired neurogenesis in the hippocampal formation (Hodge et al., 2012). As these IPCs are hypothesized to play a role in neocortical expansion, they may also be involved in the lineage-specific hippocampal expansion seen in humans.

Many of the differences between humans and chimpanzees are due to diverging gene regulatory sequences (Agoglia et al., 2021; Enard et al., 2002; Gokhman et al., 2021; King and Wilson, 1975; Wray, 2007). A recent study comparing gene expression between adult human, chimpanzee and macaque brain regions identified several genes specifically upregulated in the human hippocampus (Sousa et al., 2017). However, transcriptomic differences between primate species during specific timepoints of hippocampal development have not been investigated.

As samples of developing human and chimpanzee brain tissue are extremely limited, Induced Pluripotent Stem Cells (iPSCs) are an ideal system to conduct comparative studies of human and chimpanzee hippocampal development. Previous studies have employed iPSC-derived cortical organoids and iPSC-derived neuronal progenitor cells (NPCs) from human and chimpanzee for comparative and developmental genomic purposes (Marchetto et al., 2019; Mora-Bermúdez et al., 2016). Here, we leverage human and chimpanzee iPSC-derived hippocampal progenitors as models for comparative developmental and genomic studies with the goal of identifying species-specific differences in gene regulation during hippocampal development.

Several recent papers have recently demonstrated that transposable elements (TEs) can alter existing regulatory elements or generate entirely novel ones, as well as expand in a species- or lineage-specific manner (reviewed in Sundaram and Wysocka, 2020). Species-specific TE expansion and co-option into gene regulatory networks have been demonstrated as a mechanism for evolutionary change (Chuong et al., 2016; Fuentes et al., 2018; Jacques et al., 2013; Lynch et al., 2011; Lynch et al., 2015; Miao et al., 2020; Mika et al., 2021; Pontis et al., 2019; Trizzino et al., 2017). Endogenous Retroviruses (ERVs) and SINE-Vntr-Alus (SVAs) are among the TE families more frequently associated with gene regulatory activity in the human genome (Chuong et al., 2016; Fuentes et al., 2018; Pontis et al., 2019; Trizzino et al., 2017; Trizzino et al., 2018). SVAs encompass six subfamilies, denoted as SVA-A through -F. Of the ∼3,000 SVA copies in the human genome, nearly half are human specific, including all the SVA-E and SVA-F (Quinn and Bubb, 2014; Wang et al., 2005). The remaining half are also found in other great apes. SVAs are still replication competent and thus able to transpose in the human genome. ERVs are retrotransposons belonging to the Long Terminal Repeat (LTR) group. They are remnants of past retroviral infection events, and make up ∼8% of the human genome (Tokuyama et al., 2018). Both SVAs and ERVs were recently found to be enriched within the sequences of active cis-regulatory elements (enhancers, promoters) in hippocampal tissue as compared to other human brain regions where they are predominantly repressed (Trizzino et al., 2018). Therefore, we hypothesize that species-specific ERV and SVA transposon activity may influence the gene regulatory networks necessary for human and chimpanzee hippocampal development.

Given the key function that intermediate progenitors played in the evolution of the primate brain (Florio and Huttner, 2014; Martínez-Cerdeño et al., 2006), we sought to identify molecular differences between iPSC-derived human and chimpanzee hippocampal intermediate progenitors (hpIPCs) in terms of gene expression and the regulatory activity of non-coding regions. We specifically examined gene expression (RNA-seq and scRNA-seq), gene regulation (ATAC-seq) and functional TE activity via CRISPR-interference.

We leveraged singe-cell RNA-seq to examine the temporal trajectory of the hpIPCs during differentiation in both species. After confirming that the hpIPC differentiated cells express the appropriate neurodevelopmental markers, we conducted a transcriptomic comparison between human and chimpanzee hpIPCs. This analysis revealed over 2,500 differentially expressed genes. We then used ATAC-seq to conduct extensive analyses of differential chromatin accessibility between human and chimpanzee hpIPCs. In both species, differentially accessible chromatin regions were more likely than expected to overlap a TE insertion. Further, these regions were found as both enriched and depleted for specific TE families. Notably, species-specific enrichment for ERV and SVA sequences within differentially accessible genomic sites correlated with species-specific changes in nearby gene expression. This is likely driven by transcription factors binding to the TE-derived regulatory sequences, as we demonstrate for TBR2 and SVA-derived enhancers. Finally, we used CRISPR-interference to repress all the accessible SVAs in progenitor-like cells and demonstrated that such repression results in global changes in gene expression and affects hundreds of important neurodevelopmental genes.

This work demonstrates that two young TE families have contributed significantly to the gene regulatory differences between human and chimpanzee hippocampal development, providing insight into how the human hippocampus evolved both its unique cognitive capacity and its susceptibility to neurodegenerative disease.

## Results

### An iPSC-derived model for human and chimpanzee hippocampal intermediate progenitors

We modeled hpIPCs in humans and chimpanzees using three human and three chimpanzee iPSC lines. All six iPSC lines were validated as pluripotent in previous studies (Gallego Romero et al., 2015; Pagliaroli et al., 2021; Pashos et al., 2017; Ward et al., 2018; Yang et al., 2015; Zhang et al., 2015). For both human and chimpanzee iPSCs, we used two female cell lines and one male line.

We used a previously published method to generate hpIPCs from iPSCs (Yu et al., 2014). In this protocol, the stem cells are treated with a media containing anticaudalizing factors and a Sonic Hedgehog antagonist (DKK1, Nogging, SB431542) to generate forebrain progenitor cell types (Yu et al., 2014). It should be noted that the hpIPCs are distinct from a more general neuronal progenitor cell (pan-NPC) as the pan-NPC media is supplemented only with FGF2 and B27. Moreover, and the hpIPCs can be further induced to generate mature hippocampal CA3 pyramidal or dentate gyrus granule neurons (Sarkar et al., 2018; Yu et al., 2014).

As we were specifically interested in TBR2-positive intermediate hippocampal progenitors, we differentiated one human male and one chimpanzee male iPSC line. To ensure that the differentiation process proceeded similarly in each species, we compared the transcriptomes by conducting single-cell RNA-seq (hereafter scRNA-seq) at 24 hour intervals from the iPSC stage (day 0) to the hpIPC (day 5). We assayed a total of 18935 cells, with 4540 chimpanzee cells and 14395 human cells. These data demonstrate that the differentiation follows a similar trajectory in both species (Figure 1A). Despite the difference in final cell number between each species (Supplemental Figure 1A), all six time points overlap closely between species (Supplemental Figure 1B, 1C). Chimpanzee cells from day 2 and day 3 cells overlap more than expected with human cells from day 3 and day 4, respectively, suggesting that the chimpanzee cells are farther along in the hpIPC differentiation at these time points. However, both species align closely again by day 5, which is the time point chosen for all the genomic analyses conducted in the present study.

**Figure 1.**
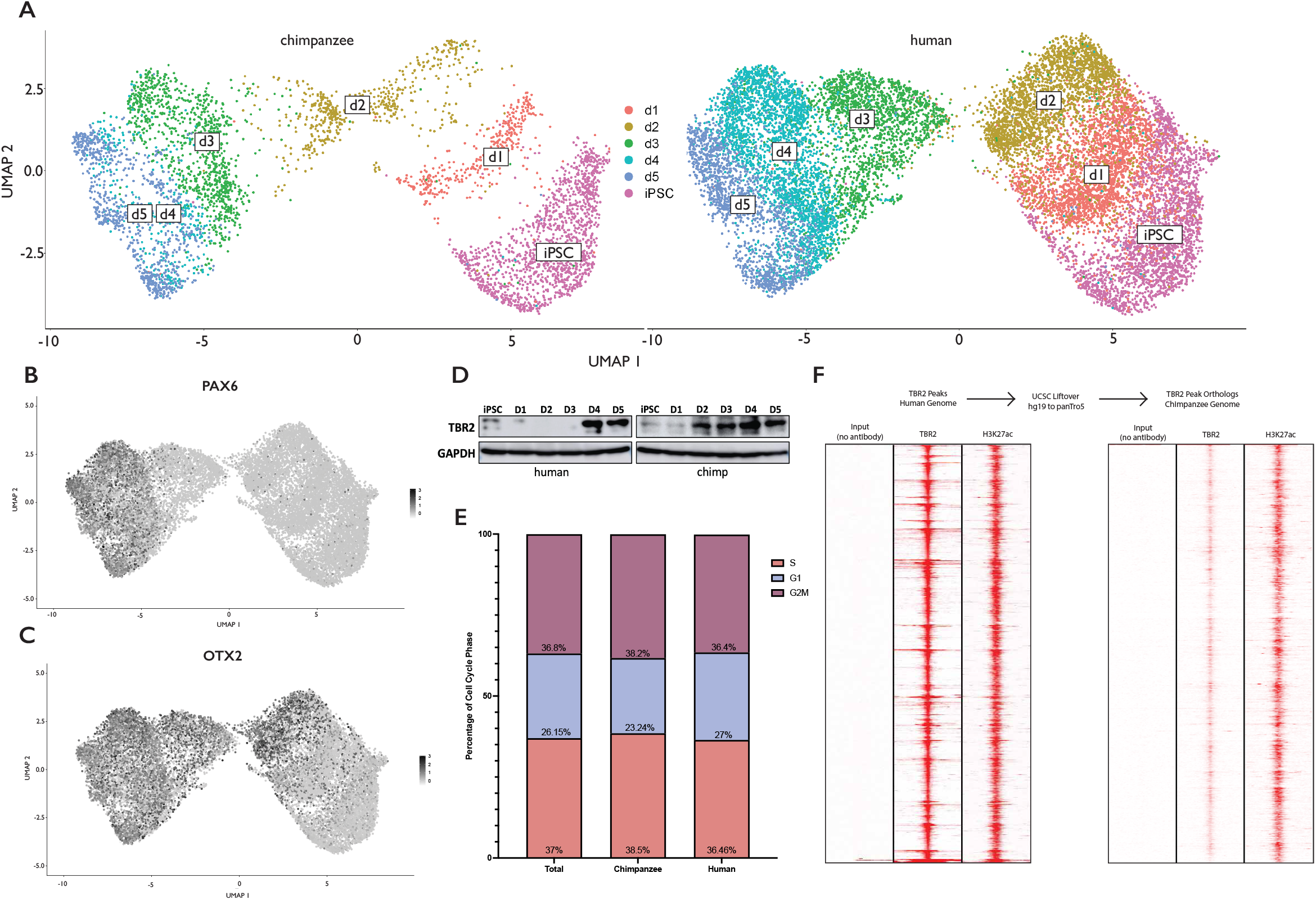
Differentiation of human and chimpanzee iPSCs into hippocampal intermediate progenitors (A) single-cell RNA-seq data of human and chimpanzee cells from iPSC to day 5 of hpIPC differentiation, grouped by time point and split by species. (B, C) PAX6 and OTX2 expression across the hpIPC differentiation time points. Darker purple indicates greater expression. (D) Western Blot of human and chimpanzee differentiated iPSCs against TBR2 from day 0 to 5 of differentiation. GAPDH is shown as the loading control. (E) Quantification of cells in S, G1 and G2M phases of the cell cycle from single-cell RNA-seq data. (F) Heatmaps of ChIP-seq against TBR2 and H3K27ac in day 5 hpIPCs from human and chimpanzee. Red indicates signal.

We further leveraged the scRNA-seq data to examine the expression of known neurodevelopmental markers and saw a progressive increase in the expression of both PAX6 and OTX2 during the 5 day differentiation (Figure 1B, 1C), which remains consistent between species (Supplemental Figure 1D, 1E). To investigate any noticeable changes to cell cycle or proliferation between species, we quantified the number of at the G1, S, and G2M phases of the cell cycle within the scRNA-seq data set, based on a set of known cell cycle genes and did not detect significant differences (Figure 1E).

To ensure that the differentiation resulted in hpIPCs specifically, we conducted a Western Blot in both species and confirmed that TBR2 expression in human cells is greatest at days 4 and 5 (Figure 1D), similarly to OTX2 and PAX6 expression (Figure 1B, 1C). In chimpanzee cells, TBR2 is expressed as early as day 2, corroborating evidence from the scRNA-seq that chimpanzee differentiation initially proceeds faster. However, we do not see evidence that TBR2 expression in chimpanzee cells is greater than TBR2 expression at day 5 in human cells. As OTX2 labels neuronal progenitors and TBR2 and PAX6 mark the intermediate progenitor cell type (Florio and Huttner, 2014, Cipriani et al., 2016, Hevner, 2019), the data demonstrate that the differentiation was successful and comparable between species. Lastly, we conducted ChIP-seq against TBR2 and H3K27ac to determine whether TBR2 was bound at the same genomic sites in both species (Figure 1G). We identified 3789 TBR2-bound regions in human hpIPCs and observed that nearly all of them also exhibited the active histone mark H3K27ac. Upon translating the coordinates of these regions to the chimpanzee genome, we observed that the 3781 orthologous regions were bound by TBR2 in chimpanzee hpIPCs. Together, these data indicate that our iPSC-derived model is suitable to study gene regulation within TBR2-positive hippocampal intermediate progenitors in both human and chimpanzee.

### Important neurodevelopmental genes are differentially expressed between human and chimpanzee hippocampal intermediate progenitors

To investigate the differences between human and chimpanzee hippocampal intermediate progenitors, we first aimed to characterize differential gene expression between the hpIPCs of the two species. After five days of treatment with the hpIPC differentiation media, we collected cells for RNA extraction (Fig. 2A). To ensure statistical power, we conducted bulk RNA-seq on two replicates each of all six cell lines (i.e. three biological replicates and six technical replicates per species). The differentiation, the harvesting, and the RNA processing were performed in mixed batches with samples from both species to prevent batch effects. The libraries were sequenced using Illumina NextSeq500, generating 100 bp Paired-End reads. Non-orthologous genes were omitted from the analysis and a total of 2,588 genes were identified as differentially expressed (FDR <0.05 and log2foldChange > 1.5 or < -1.5). The genes with log2foldChange > 1.5 were more highly expressed in the human hpIPCs (“Human UP”) while the genes with log2foldChange < -1.5 were more highly expressed in the chimpanzee hpIPCs (“Chimp UP”). In total, 1,686 (65.1%) of the differentially expressed genes (DEG) were “Human UP” and 901 (34.9%) of the DEG were “Chimp UP” genes (Fig. 2B, Supplementary Table 1.1).

**Figure 2.**
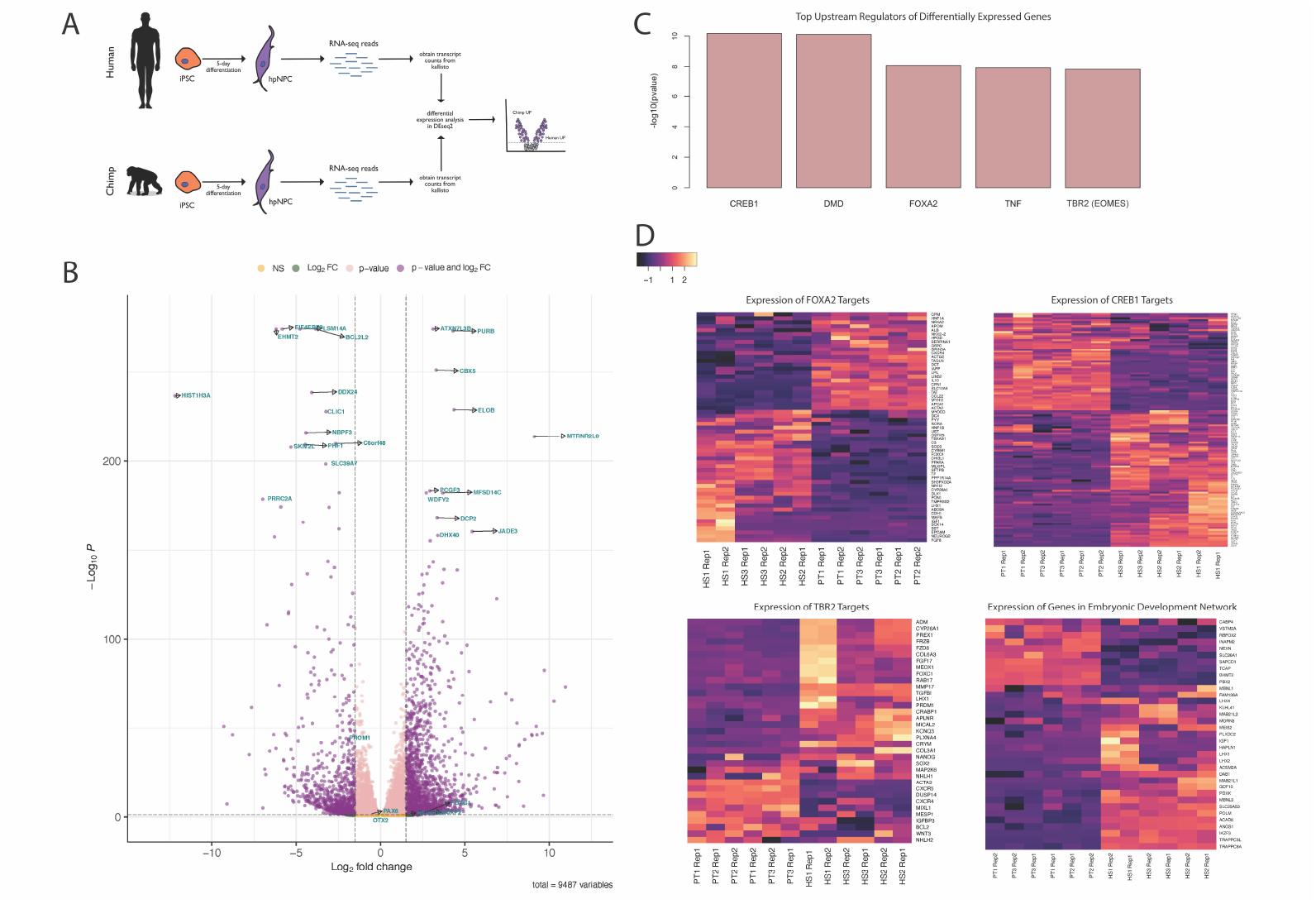
Differential Gene Expression In Human and Chimpanzee hpIPCs. (A) Schematic of iPSC differentiation followed by RNA-seq library generation and analysis. (B) Volcano plot depicting “Human UP” genes (right) and “Chimp UP” genes (left) with a log2FoldChange threshold of 1.5 and -1.5, respectively, and a p-value threshold of 0.05. (C) Top upstream regulators of the differentially expressed genes predicted by Ingenuity Pathway Analysis, ranked by -log10(p-value). (D) Heatmaps depicting the expression of genes predicted to be under the control of the transcription factors CREB1, FOXA2 and TBR2 or predicted to play a role in embryonic development. The rows of the heatmaps indicate the transcript names, columns indicate the species, sample number and replicate. “HS1 Rep 1” indicates Homo sapiens sample 1 replicate 1 and “PT2 Rep 2” indicates Pan troglodytes sample 2 replicate 2

Hippocampal neurodevelopmental markers *PAX6, OTX2, NEUROD1* and *FOXG1* were found as highly expressed in both species. This is consistent with the immunostaining in demonstrating that the hpIPCs differentiation was successful and that the progenitors from both species are comparable.

The “Human UP” genes include *FOXP2, MTRNR2L8, DHX40, VPS13B, WDFY2*, and *PURB*, all of which are associated with neurodevelopment or neurodegenerative disease (Abrajano et al., 2009; Hickey et al., 2019; Kamboh et al., 2019; Kolehmainen et al., 2003; MacDermot et al., 2005; Mathys et al., 2019; Sin et al., 2015; Taher et al., 2014). The “Chimp UP” genes include *HIST1H3A, BLC2L2*, and *CLIC1* which have been reported as expressed in the hippocampus and associated with sleep, AD, and neurite outgrowth (Averaimo et al., 2014; Datson et al., 2009; Wei, 2020).

To understand the transcriptional programs driving these differences in gene expression, we conducted an Ingenuity Pathway Analysis. Three of the top five upstream regulators predicted by the pathway analysis were the transcription factors CREB1, FOXA2 and TBR2 (EOMES) (Fig. 2C). CREB1 is known to regulate genes involved in the nervous system and neurodevelopment (reviewed in Sakamoto et al., 2011), FOXA2 controls dopaminergic neuronal development and disease (Kittappa et al., 2007), and TBR2 plays a crucial role in cortical and hippocampal neurogenesis and is the signature marker of the intermediate progenitor population (Cipriani et al., 2016; Englund, 2005). This pathway analysis was consistent with the RNA-seq data as predicted targets of all three transcription factors were also found to be among the 2,588 genes differentially expressed between human and chimpanzee hpIPCs (Fig. 2D, Supplementary Table 1.2-1.4). The pathway analysis also determined that several of the differentially expressed genes are involved in embryonic development (Fig. 2D, Supplementary Table 1.5). Overall, these findings indicate that previously characterized neurodevelopmental gene regulatory networks are utilized differently during human and chimpanzee hippocampal development.

### Human-specific chromatin accessibility patterns in hippocampal intermediate progenitors

After identifying differentially expressed genes and the transcriptional networks that may be involved, we sought to identify cis-regulatory differences between the human and chimpanzee hpIPCs. To this end, we conducted ATAC-seq on the hpIPCs from both species. We used the same batches of differentiated iPSCs for the ATAC-seq as we did for the RNA-seq (i.e. from the same batch of differentiation), and generated 100 bp long Paired-End Illumina reads.

We first performed a “human-centric” analysis. We aligned the ATAC-seq reads from all six cell lines to the respective reference genome assemblies (hg19 for the human cell lines, panTro5 for the chimpanzee cell lines) and only retained uniquely mapped reads with high mapping quality (Samtools Q = 10 filtering). Next, we identified regions of accessible chromatin (peaks; FDR < 0.05) in all three human cell lines. Only peaks replicated in all the three human lines were retained. To carry out a proper comparison, we only retained replicated human ATAC-seq peaks with orthologs in the chimpanzee genome (see methods; Fig. 3A). This filtering ultimately resulted in 82,235 human ATAC-seq peaks replicated in all the human cell lines and with orthologs in the chimpanzee genome. These 82,235 regions were used for downstream analysis. We found that the chromatin accessibility at these regions was highly reproducible across all three human cell lines (Fig. 3B).

**Figure 3.**
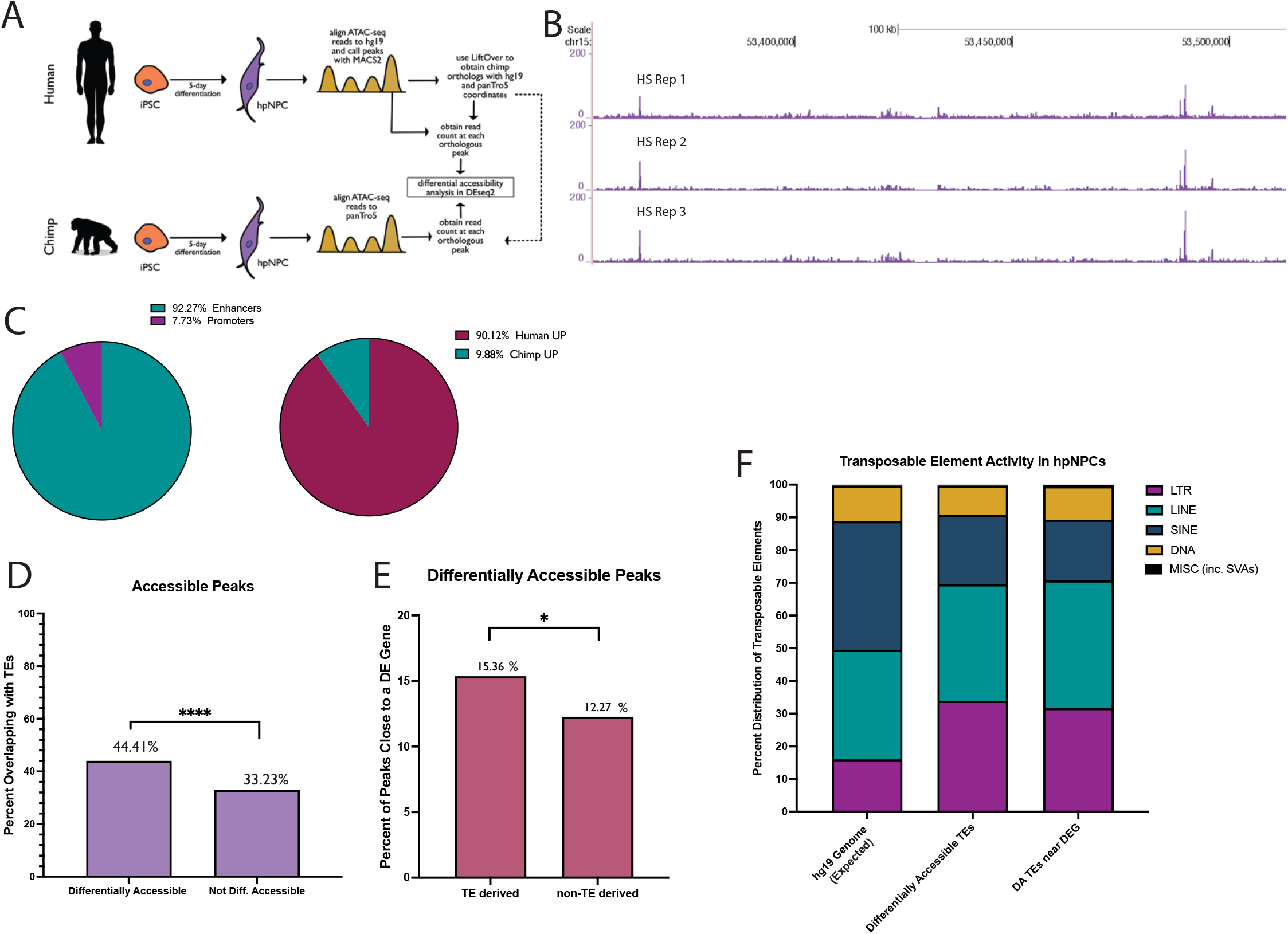
Human Centric Chromatin Accessibility Analysis. (A) Schematic of iPSC differentiation followed by ATAC-seq library generation and analysis. (B) UCSC Genome Browser visualization of human ATAC-seq libraries (HS Rep 1, Rep 2, Rep 3) aligned to hg19 genome assembly. (C) Distribution of the differentially accessible (DA) peaks into enhancers or promoters, and into Human UP (greater accessibility in humans) or Chimp UP (greater accessibility in chimpanzees). (D) Differentially accessible chromatin regions (p < 0.05, n = 3006) are more likely to overlap with transposable elements (TE) than non-differentially accessible regions (p > 0.9, n = 3,006). (E) The 1335 TE-derived DA regions are more likely to be near a differentially expressed gene, as compared to DA regions that do not overlap a TE. (F) Distribution of the five major TE classes in the human genome, compared with their distribution among the TE-derived DA chromatin regions, and among the TE-derived DA chromatin regions proximal to a differentially expressed gene

Next, we quantified the ATAC-seq read depth for each of the 82,235 regions and used DESeq2 to identify sites exhibiting differential chromatin accessibility between the two species. In total, we identified 3,006 differentially accessible regions (FDR <0.05; log2foldChange > 1.5 or < -1.5; Supplementary Table 2.1-2.2). Of these regions, 92.3% were located at least 1 kb away from the closest transcription start site (TSS), suggesting they could be putative enhancers, while the remaining were putative promoters (Fig. 3C). As expected, given that this analysis was performed with a human-centric approach, 90.1% of the differentially accessible (DA) peaks were significantly more accessible in the human hpIPCs relative to chimpanzee hpIPCs (“Human UP”; Fig. 3C).

As TE insertions can be a source of cis-regulatory evolution, we examined whether the DA regions were more likely to overlap with a TE than those which were accessible to the same degree in both species (non-DA, Fig. 3D and Supplementary Table 2.4-2.5). We observed that 1,335/3,006 (44.4%) DA regions overlapped a TE (Supplementary Table 2.3). This is significantly higher than what was observed for the non-DA peaks (33.2% overlapped a TE; Two-sided Fisher’s Exact Test, p < 0.0001; Fig. 3D). This indicates that chromatin regions with human-specific accessibility are significantly more likely to be TE-derived than the regions with accessibility levels conserved between human and chimpanzee.

Next, we associated the nearest gene to each DA region and found that TE-derived DA regions were significantly more likely to be near a differentially expressed gene relative to non-TE derived DA regions (Two-sided Fisher’s Exact Test, p = 0.016; Fig. 3E). Finally, we investigated whether specific TE families were overrepresented among the TE-derived DA regions and found enrichment for LTRs. While LTRs account for ∼16% of all human annotated TEs, they represented 33.9% of the TEs overlapping DA regions in our human-centric analysis (Two-Sided Fisher’s Exact Test p < 0.0001; Fig. 3F and Supplementary Table 2.6). Of these enriched LTRs, 97.1% were ERVs. Notably, 31.7% of those LTR-derived DA regions were located near a differentially expressed gene (Two-Sided Fisher’s Exact Test p < 0.0001).

Together, these data indicate that there are TE-derived cis-regulatory elements that have significantly greater accessibility in humans than chimpanzees during hippocampal neurogenesis. These TE insertions preceded the human-chimpanzee split, but the difference in accessibility is species-specific, suggesting that the co-option into gene regulatory networks took place after the human-chimpanzee divergence.

### Chimpanzee-specific chromatin accessibility patterns in hippocampal intermediate progenitors

We repeated the ATAC-seq analysis as described above, but this time with a “chimpanzee-centric” approach. We started from a set of 72,211 peaks found as replicated in all the three chimpanzee lines and with orthologs in both species (Figs. 4A,B). With this approach, we identified 3,806 ATAC-seq peaks as differentially accessible (DA) between human and chimpanzee (FDR <0.05; log2foldChange > 1.5 or < - 1.5; Supplementary Table 3.1-3.2), 82% of which displayed greater accessibility in chimpanzee as compared to humans (i.e. “Chimp UP”; Fig. 4C). Similar to what we observed with the human-centric analysis, 97.9% of the 3,806 DA regions were putative enhancers (distance > 1Kb from TSS; Fig. 4C).

**Figure 4.**
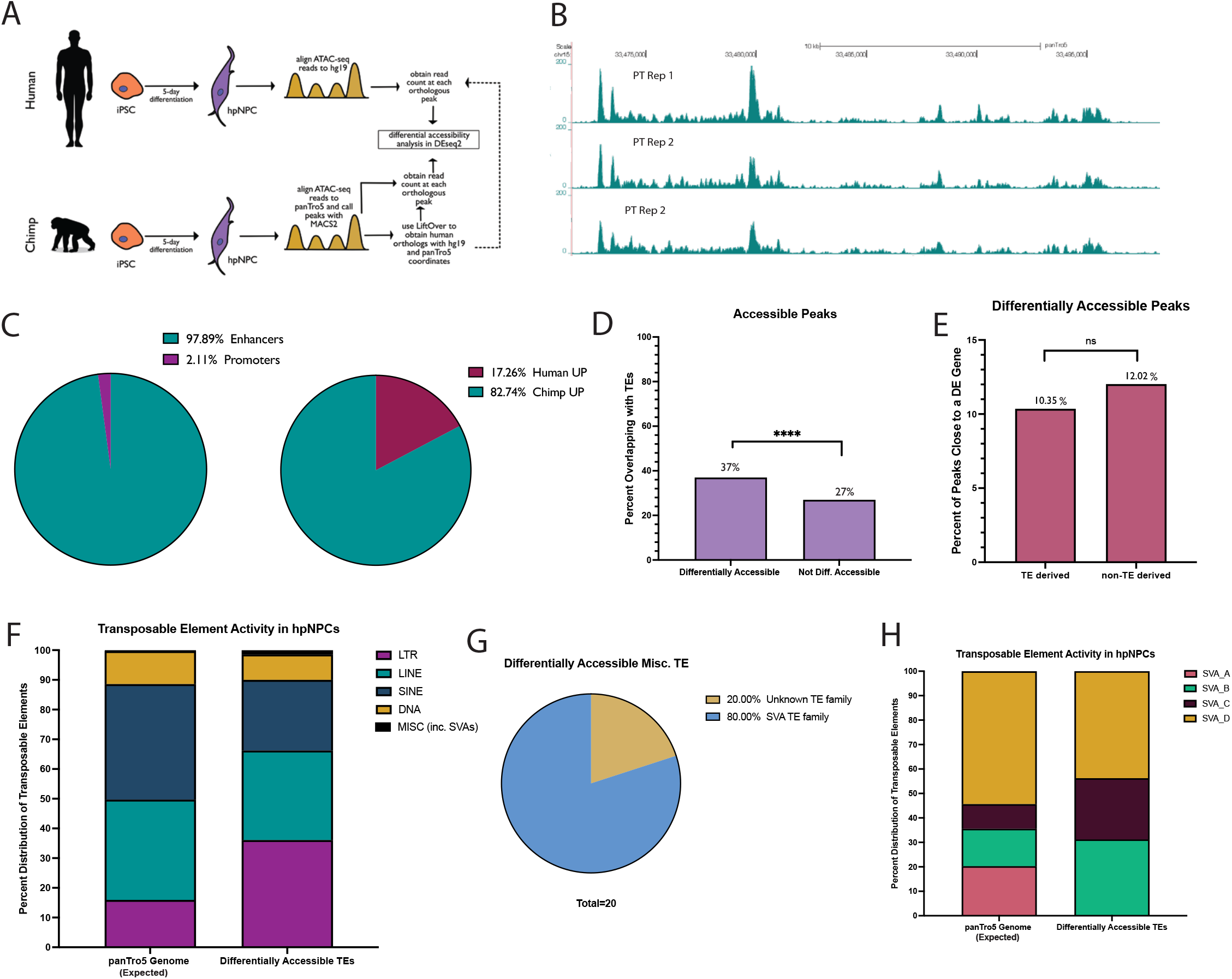
Chimpanzee Centric Chromatin Accessibility Analysis. (A) Schematic of iPSC differentiation followed by ATAC-seq library generation and analysis. (B) UCSC Genome Browser visualization of human ATAC-seq libraries (PT Rep 1, Rep 2, Rep 3) aligned to hg19 genome assembly. (C) Distribution of the differentially accessible (DA) peaks into enhancers or promoters, and into Human UP (greater accessibility in humans) or Chimp UP (greater accessibility in chimpanzees). (D) Differentially accessible chromatin regions (p < 0.05, n = 3,806) are more likely to overlap with transposable elements than other accessible regions (p > 0.9, n = 3806. (E) Distribution of DA regions that overlap a TE and proximity to genes differentially expressed between human and chimpanzee hpIPCs. (F) Distribution of the five major TE classes in the chimpanzee genome, compared with their distribution among the 1410 TE-derived DA chromatin regions (G) Breakdown of the 20 Miscellanous TEs represented in the 1,410 TE-derived DA chromatin regions. (H) Distribution of SVA family of TEs in the human genome as compared to their distribution in the TE-derived DA chromatin regions (n = 16).

As seen in the human-centric analysis, chimpanzee-specific DA peaks were more likely to be TE-derived than those similarly accessible across species (Two-sided Fisher’s Exact Test, p < 0.0001; Fig. 4D and Supplementary Tables 3.3-3.5). Of the chimpanzee DA regions, 37.1% overlapped an annotated chimpanzee TE, compared to only 28.1% of the non-DA regions (Two-sided Fisher’s Exact Test, p < 0.0001; Fig. 4D). However, the TE-derived enhancers in this chimpanzee-centric analysis were no more likely than the non-TE derived ones to be located near differentially expressed genes (Fig. 4E).

We found enrichment for LTRs, which account for approximately 16% of the chimpanzee TEs but represented 36.1% of the TE-derived DA regions (Two-sided Fisher’s Exact Test, p-value < 0.0001; Fig. 4F and Supplementary Table 3.6; 98.2% were ERVs), and SVAs, which account for just 0.25% of annotated chimpanzee TEs but represented 1.1% of the TE-derived DA regions (Two-sided Fisher’s Exact Test, p < 0.0001, Fig 4G and Supplementary Table 3.8). In particular, the SVA-B and C subfamilies were the most enriched (Fig. 4H). Together, these data indicate that chimpanzee ERV and SVA transposons were co-opted into regulatory elements important for the developing chimpanzee hippocampal intermediate progenitors.

### Genomic features underlying species-specific LTR enrichment at hpIPC enhancers

We aimed to further investigate genomic features potentially underlying the LTR enrichment among the differentially accessible hippocampal progenitor regions. To this end, we conducted a motif analysis using the MEME suite (Bailey et al., 2015). Binding motifs for CTCF and EGR2 were detected as enriched in the LTR-derived differentially accessible regions identified from both the human-centric and chimpanzee-centric analyses (Figs. 5A, 5C). EGR2 is an early response gene involved in learning and memory, in the brain’s response to stimuli, and in hippocampal synaptic plasticity (Cheval et al., 2012; Mukherjee et al., 2021; Poirier et al., 2007). CTCF, a well-known regulator of chromatin structure, has been implicated in various neurodevelopmental disorders (reviewed in Davis et al., 2018)

**Figure 5.**
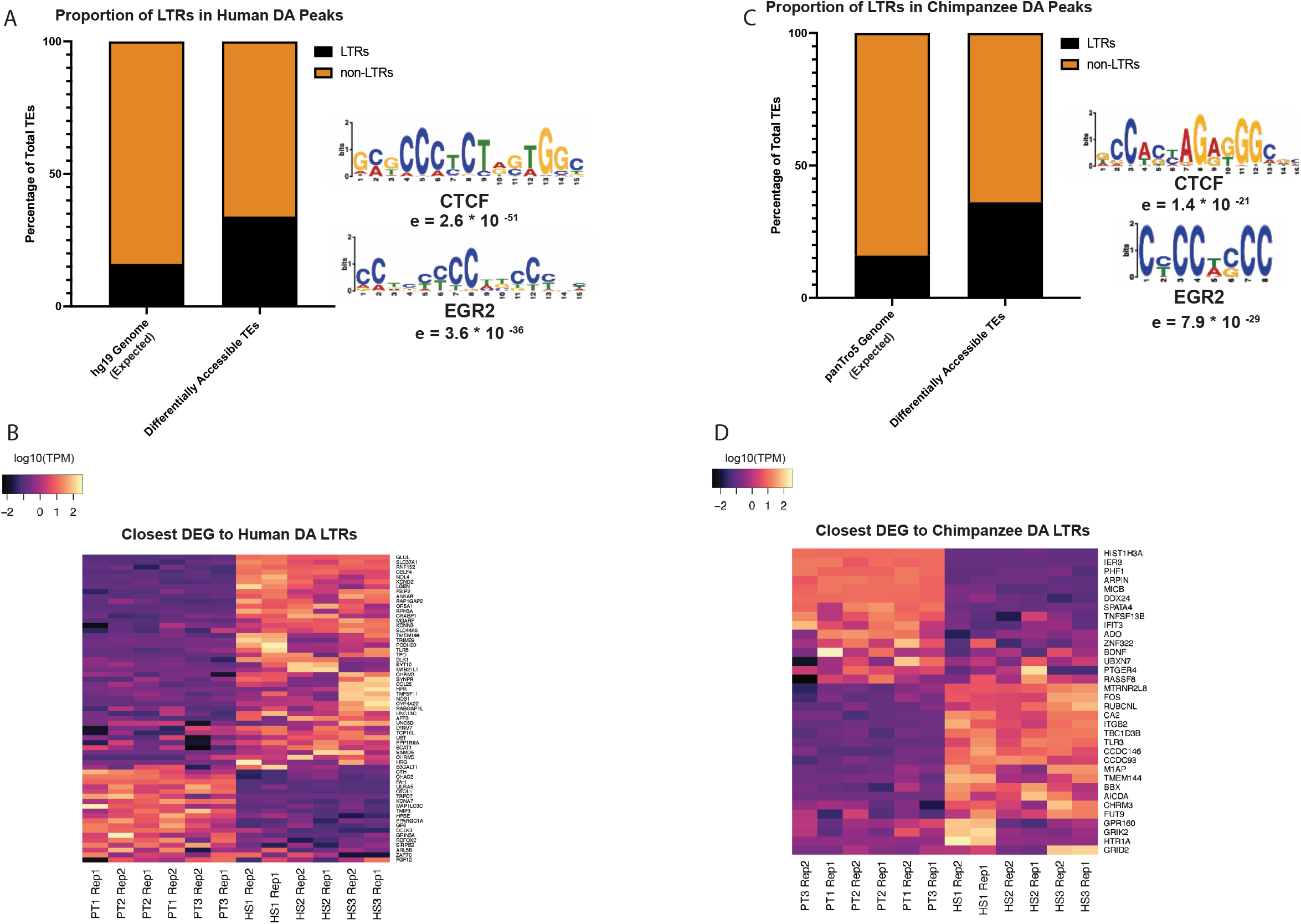
LTRs Are Enriched Among Human and Chimpanzee Differentially Accessible Transposons. (A) Distribution of LTRs compared to non-LTRs in the human genome and in the 1,335 human TE-derived DA regions, with predicted binding motifs. (B) Differentially expressed genes close to the human-enriched DA LTRs. (C) Distribution of LTRs compared to non-LTRs in the chimpanzee genome and in the 1,410 chimpanzee TE-derived DA regions, with predicted binding motifs. (D) Differentially expressed genes close to the chimpanzee-enriched DA LTRs.

We identified 63 differentially expressed genes located near the human-enriched LTRs (Fig. 5B and Supplementary Table 2.7). These included *GLUL*, a glutamine synthetase hypothesized to provide neuroprotection in AD patients, (Kohane and Wood, 2021) and *DLK1*, a Notch ligand involved in SVZ neurogenesis (Ferrón et al., 2011), both of which are upregulated in human hpIPCs compared to chimpanzee hpIPCs.

We identified 34 differentially expressed genes located near the chimp-enriched LTRs (Fig. 5D and Supplementary Table 3.7). For the chimp-centric analysis the differentially expressed genes located near TE-derived enhancers with species-specific accessibility included *HIST1H3A* and *MTRNR2L8* (Fig. 5D). These genes were highly upregulated in the chimpanzee and human hpIPCs, respectively (Fig. 5D). Importantly, *HIST1H3A* has been associated with autism spectrum disorders and sleep deprivation (Crawley et al., 2016; Wei, 2020) while *MTRN2L8* has been reported as upregulated in Alzheimer’s patients (Mathys et al., 2019).

### Human-Specific SVAs play a major role in hippocampal neurogenesis

As mentioned earlier, the chimpanzee-centric analysis also identified SVA transposons as enriched within chimpanzee-specific enhancers (Fig. 6A and Supplementary Table 3.8). These SVAs were enriched for the identified binding motifs of the neurodevelopmental factors ASCL1, ZIC1, and KLF8 (Andersen et al., 2014; Aruga, 2004; Yi et al., 2014) as well as the JUN/FOS AP-1 dimer, which is a known enhancer activator (Raivich, 2008; Raivich and Behrens, 2006) (Fig 6A).

**Figure 6.**
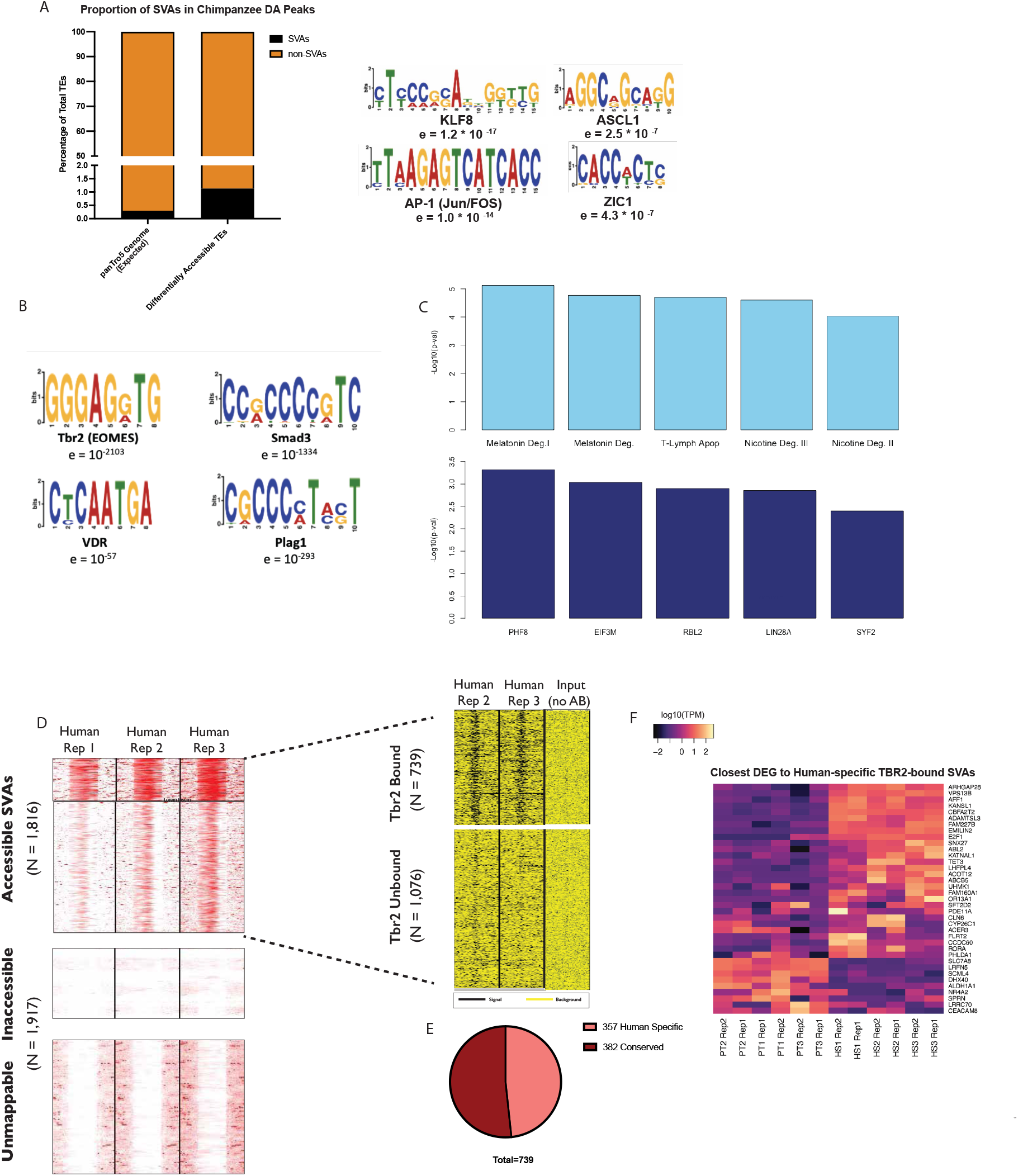
Human-specific SVAs Bind Tbr2 and Influence Neurodevelopment (A) The SVAs enriched within the differentially accessible chimpanzee peaks are enriched for binding motifs of neurodevelopmental transcription factors KLF8, ZIC1, and ASCL1 as well as the AP-1 dimer. (B) Human SVAs are enriched for the binding motif of TBR2. (C) Genes proximal to human-specific SVAs are predicted to be regulated by neurodevelopmental TFs and function in important neuronal pathways. (D) 1,816 human SVAs exhibit ATAC-seq signal in hpIPCs (red = signal, white = noise) and of these, 739 exhibit TBR2 ChIP signal (black = signal, yellow = background). (E) Breakdown of 739 accessible, TBR2 bound SVAs into “human-specific” and “conserved in chimpanzees”. (F) Differentially expressed genes close to the human-specific TBR2-bound SVAs.

It is important to remark that all the analyses shown so far were exclusively based on genomic sites with characterized orthologs in both species, to ensure an “apple-to-apple” comparison. However, nearly 2,000 SVA copies, including the entire SVA-E and F subfamilies, are exclusive to the human genome. Given that previous studies found that SVAs are highly enriched in active enhancers and promoters of the human hippocampus (Trizzino et al., 2018), we sought to investigate this further. We focused on the SVA copies exclusively present in the human genome and not in any other primate genome (hereafter human-specific SVAs). We performed sequence-based motif analysis for all the human-specific SVAs and identified the binding motif for the hpIPCs signature factor TBR2 (EOMES) as the most enriched (p = 10^×2103^; Fig. 6B). Motifs for other transcription factors associated to hippocampal neurogenesis and function were also recovered (SMAD4, VDR, PLAG1; Fig. 6B).

Next, we annotated all of the genes found within 50 kb from each human-specific SVA. A total of 2,216 genes were recovered using this approach. We performed Pathway Analysis on this set of genes and found that Melatonin Degradation and Nicotine Degradation were the two most significantly enriched pathways (p = 7.6 × 10^×6^ and p = 2.5 × 10^×5^ respectively; Fig. 6C) and PHF8 was recovered as the top upstream regulator for the gene network (Fig. 6C). Notably, both Melatonin and Nicotine Degradation pathways are strongly active in the human hippocampus. PHF8 is a histone demethylase that contributes to the regulation of mTOR. The mTOR pathway is hyperactive in the human hippocampus where it regulates the protein synthesis-dependent plastic changes underlying learning and memory (Bekinschtein et al., 2007; Fortress et al., 2013; Graber et al., 2013). Mutations in the *PHF8* gene cause cognitive impairment and intellectual disability (Chen et al., 2018).

Therefore, we sought to determine the contribution of human-specific SVAs to the TBR2-mediated gene regulatory network in human hpIPCs. Using our ATAC-seq data, we identified 1,816 human SVAs as accessible in human hpIPCs (Fig. 6D). Of these, nearly a quarter (434) displayed high accessibility, while 1,382 were moderately accessible (Fig. 6D). Next, we used chromatin immunoprecipitation followed by sequencing (ChIP-seq) to profile TBR2 binding in two human lines at day 5 of hpIPC differentiation. As with the previous sequencing experiments, we generated 100 bp long Paired-End reads and only retained uniquely mapping high quality reads (Samtools q=10 filtering) in order to maximize our chance to properly map reads on repetitive regions. This experiment revealed that 739 of the accessible SVAs showed TBR2 signal in the two human lines (Fig. 6D and Supplementary Table 2.8). Notably, 257 of the (48.3%) TBR2-bound SVAs were human-specific. (Fig. 6E and Supplementary Table 2.9). TBR2-bound human-specific SVAs were located near 37 genes that our RNA-seq analysis identified as differentially expressed between human and chimpanzee (Fig. 6F and Supplementary Table 2.10). These genes include *VPS13B* (up in humans), which is responsible for a rare developmental disease known as Cohen Syndrome (Kolehmainen et al., 2003), *NR4A2* (down in humans), which has been implicated in neurodevelopmental language impairment (Reuter et al., 2017), *DHX40* (down in humans), which is implicated in Alzheimer’s (Taher et al., 2014), and *E2F1* (up in humans), which is a cell cycle regulator associated with several neurodegenerative diseases such as Alzheimer’s (Zhang et al., 2010).

In summary, these data support a model in which human-specific SVAs provided a substrate for binding sites of TBR2 and other important hippocampal regulators. Therefore, it is likely they were co-opted in the gene regulatory networks active during hippocampal neurogenesis, which led to human-specific regulation of key genes.

### CRISPR-mediated SVA repression has massive repercussions on global gene expression

To further assess the contribution of SVA transposons to the gene regulation of hippocampal intermediate progenitors, we leveraged CRISPR-interference to simultaneously repress most of the active SVAs. We utilized NCCIT cells treated with retinoic acid (RA) as the experimental system for this purpose. The NCCIT cell line is derived from embryonal carcinoma and thus exhibits a gene expression signature highly similar to human embryonic stem cells (Fuentes et al., 2018; Barnada et al., 2022). Importantly, NCCITs treated for 7 days with RA differentiate into intermediate neural progenitor-like cells expressing both PAX6 and TBR2 (Mandal et al., 2015; Fig 7A). RNA-seq tpms suggest very strong correlation between RA-treated NCCITs and human day-5 hpIPCs (Pearson correlation p < 2.2 × 10^×16^, R=0.99). We cloned a stable NCCIT line with a permanently incorporated doxycycline-inducible, catalytically-dead Cas9 fused to a repressive KRAB domain (dCas9-KRAB). The KRAB domain deposits repressive histone methylation (H3K9me3) to the regions targeted by the dCas9 via single guide-RNAs (sgRNAs). Into this same line, we also permanently knocked-in two single guide-RNAs (sgRNAs) that are able to target >80% of all SVAs (Pontis et al. 2019). Hereafter, we refer to the RA-treated CRISPR line as RA-NCCITs. Remarkably, exposing the RA-NCCITs to doxycycline for 72 hours was sufficient to induce dCas9 activation (Fig. 7B) and the deposition of repressive histone methylation (H3K9me3) on over 2,500 previously unmethylated SVAs (Fig. 7C). We next performed RNA-seq on the RA-NCCITs with or without doxycycline treatment (three replicates per condition). First, we observed that the genome-wide expression levels (TPMs) of the RA-NCCITs where highly correlated with those of the iPSC-derived human hpIPCs (Pearson correlation = 0.93; p < 2.2 × 10^×16^). This indicates that RA-treated NCCITs are appropriate to model hippocampal intermediate progenitors.

**Figure 7.**
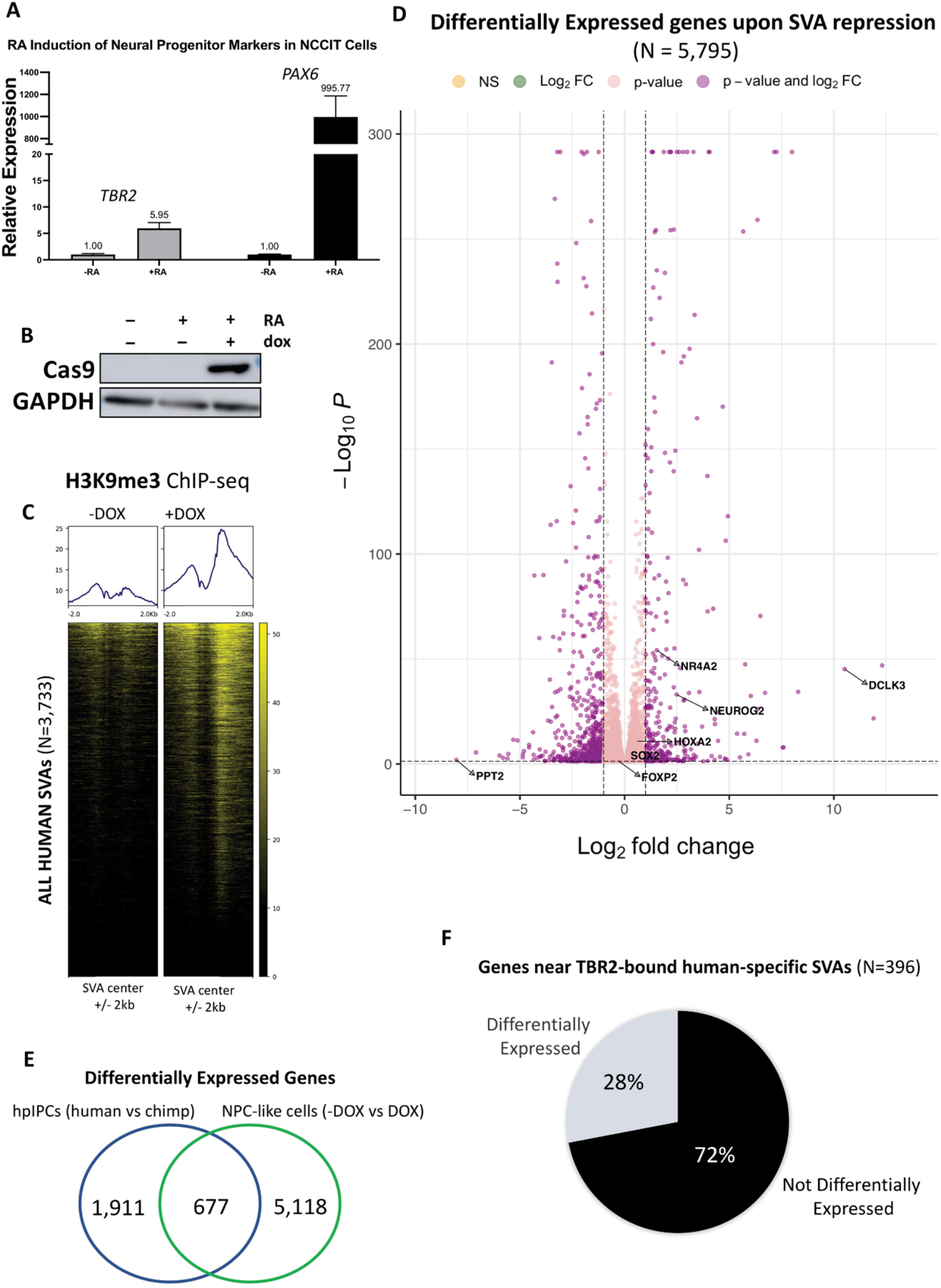
CRISPR-interference on human SVAs. (A) Treating NCCITs with Retinoic Acid for 7 days leads to induction of TBR2 and PAX6, as shown in qRT-PCR. (B) Treating the RA-NCCIT stable CRISPR line for 72 hours with doxycycline leads to dCas9 activation, as shown in the Cas9 immunoblot. (C) dCas9 activation results in deposition of H3K9me3 at most human SVAs as shown in the H3K9me3 ChIP-seq heatmap. (D) Volcano plot depicting genes differentially expressed upon doxycycline treatment. (E) Venn diagram showing overlap between genes differentially expressed between human and chimpanzee’s hippocampal intermediate progenitors and between RA-NCCITs with and without doxycycline treatment (i.e. with and without SVA repression). (F) Pie chart illustrating the fraction of TBR2-controlled genes that are differentially expressed upon SVA repression in RA-NCCITs.

Then, we compared the expression levels of the RA-NCCITs with or without doxycycline treatment and identified 5,795 differentially expressed genes (FDR < 0.05; Fig. 7D and Supplementary Table 4.1). Of these genes, 677 were previously identified as differentially expressed when comparing human to chimpanzee hpIPCs (Fig. 7E; Supplementary Table 4.2). In other words, the expression of over a quarter (26.1%) of the genes that exhibited human-specific expression signature in hippocampal intermediate progenitors seem to be under control of SVA transposons. Importantly, one of these genes was *FOXP2*, which has been associated with the evolution of language and is implicated in several speech-disorders (Enard, 2011; Liégeois et al., 2016; MacDermot et al., 2005).

Since the gRNAs used for this experiment were originally designed to target a DNA sequence shared by the SVAs with the LTR5H family (Pontis et al. 2019), we restricted the analysis to the genes that are associated to human SVAs (i.e. only considering the genes that represent the closest gene to an annotated SVA: hereafter SVA-genes). By doing so, we found 611 SVA-genes as differentially expressed in RA-NCCITs upon SVA repression (Supplementary Table 4.3). Of these, 90 were previously identified as differentially expressed when comparing human to chimpanzee hpIPCs (Supplementary Table 4.4). Thus, the expression of these 90 genes can be bona-fide considered as directly regulated by SVA-derived enhancers in both primary hippocampal progenitors and in the NCCIT cell line. Remarkably, the large majority of these genes (72.5%) had decreased expression upon SVA repression. These include *SOX2, FGF2, PRDM1, NTRK2*, and *TFAP2B* (Supplementary Table 4.4).

Finally, we examined the genes previously identified as being near a TBR2-bound human-specific SVA in iPSC-derived hpIPCs, and found that nearly a third of them lose expression in RA-NCCITs upon SVA repression (Fig. 7F). In summary, our functional experiments indicate a widespread role for human-specific SVA transposons as cis-regulatory elements during hippocampal neurogenesis.

## Discussion

The hippocampus is susceptible to specific neurodegenerative disorders such as Alzheimer’s Disease but may have also played an important role in the evolution of human cognition. Spatial memory, which is attributed to the hippocampus, may have contributed to the geographic expansion of ancient humans. Characterizing the human-specific gene regulatory networks of hippocampal development provides insight into its role in human evolution. Though there is no consensus on whether the cognitive phenotypes seen in AD are uniquely human, understanding the unique properties of the human hippocampus may lead the way to potential treatments. Thus, the work described here is relevant both in terms of evolutionary developmental biology and evolutionary medicine.

To study the evolution of the human hippocampus from a developmental standpoint, we investigated the extent to which TEs contributed to gene expression profiles of human and chimpanzee hippocampal intermediate progenitors. TEs account for nearly ∼50% of the human genome composition and many elegant studies have established that at least a fraction of the TEs can regulate the host genes in humans and other primates (Chuong et al., 2013; Chuong et al., 2016; Cosby et al., 2021; del Rosario et al., 2014; Du et al., 2016; Fuentes et al., 2018; Jacques et al., 2013; Judd et al., 2021; Lynch et al., 2011; Lynch et al., 2015; Mika et al., 2021; Modzelewski et al., 2021; Okhovat et al., 2020; Rayan et al., 2016; Schmidt et al., 2012; Sundaram et al., 2014; Trizzino et al., 2017; Trizzino et al., 2018; Ward et al., 2018; Xie et al., 2013).

The hpIPCs are a transient progenitor population during a critical developmental stage in the Sub-ventricular zone (SVZ) of both the hippocampus and neocortex (Bulfone et al., 1999; Cipriani et al., 2016; Englund, 2005; Kimura et al., 1999), and there is consensus that this progenitor population may have played a role in the evolution of brain volume in mammals (Florio and Huttner, 2014; Martínez-Cerdeño et al., 2006). To study the developmental evolution of human hpIPCs, we leveraged a comparative approach centered on differentiating human and chimpanzee iPSCs into a neuronal population which closely recapitulates hpIPCs, as demonstrated by the high expression of signature markers such as TBR2 and PAX6.

The transcriptomes of human and chimpanzee hpIPCs have not been previously compared, largely due to the limited availability of primary tissue. Here, we carried out this comparison using our iPSC-derived system and identified profound differences between the two species, with over 2,500 genes differentially expressed at this stage. These genes include several which were previously associated with cognitive function, language, neurodevelopment and neurodegeneration, many of which are upregulated in humans relative to the chimpanzee (e.g. *MTRNR2L8, DHX40, VPS13B, WDFY2, PURB*).

We demonstrate that species-specific enhancers significantly contributed to the gene expression differences we identified. This is consistent with recent studies that used primary brain tissues from several species to profile species-specific cis-regulatory activity in mammals (Emera et al., 2016; Reilly et al., 2015). Importantly, we find that these species-specific enhancers are enriched for young transposable elements. Several studies have identified evolutionarily young L1 LINEs as active in the brain during different developmental stages, suggesting that they could serve as alternative promoters for many genes involved in neuronal functions (Coufal et al., 2009; Jönsson et al., 2019; Sur et al., 2017; Thomas et al., 2012; Zhao et al., 2019). Here, we identified other young transposable elements, ERVs and SVAs, as regulators of intermediate progenitor gene expression. The identification of ERVs as candidate enhancers in human and chimpanzee intermediate progenitors is consistent with previous studies that demonstrated that ERVs heavily impact the gene regulatory programs during immune response (Chuong et al., 2013; Chuong et al., 2016), in pluripotency maintenance and development (Coluccio et al., 2018; Fuentes et al., 2018; Miao et al., 2020), in the mammalian placenta (Lynch et al., 2011; Lynch et al., 2015; Mika et al., 2021), in the primate liver (Trizzino et al., 2017), and in many cancer types (Ito et al.; Ivancevic and Chuong, 2020; Shah et al., 2021). Similar to the L1s, the ERVs also possess a well-defined cis-regulatory architecture (e.g. they have their own promoter), and this may have had a role in the co-option of these TEs as functioning cis-regulatory elements.

The SVAs are particularly interesting from a human evolution standpoint, given that half of the known copies are exclusively present in our species. Moreover, SVAs are among the few transposable elements that still exhibit active transposition in the human genome. We and others have previously revealed that SVAs can be source of enhancers in primates (Playfoot et al., 2021; Pontis et al., 2019; Trizzino et al., 2017; Trizzino et al., 2018). The repression of some SVAs by specific zinc-finger proteins at specific stages of neuronal development is also a crucial mechanism for successful neurogenesis (Playfoot et al., 2021; Pontis et al., 2019; Turelli et al., 2020). Here, we demonstrate that SVAs are pervasive regulators of hippocampal neurogenesis and they act as enhancers in the hippocampal intermediate progenitor population. By using CRISPR-interference (CRISPR-i), we show that repressing hundreds of normally “de-repressed” SVAs alters the expression of thousands of genes. Intriguingly, our CRISPR-i experiments revealed that global SVA repression leads to the attenuation of the expression levels of over a quarter of the ∼2,500 genes previously identified as with human-specific gene expression signal in hippocampal intermediate progenitors. These include critical neurodevelopmental regulators such as *FOXP2, HAND2, MEF2C, SOX2*, and *SOX4*. In normal conditions, these genes are more highly expressed in humans relative to the chimpanzee, but upon SVA repression these differences were diminished.

In conclusion, our findings indicate that the development of hippocampal neurons has been profoundly affected by the domestication of young transposable elements. These young TEs have been co-opted as functional enhancers and promoters and ultimately rewired the expression of hundreds of critically important neuro-regulators in the developing human hippocampus. The human-specific gene expression and the associated TE-derived enhancers we have identified here may play important roles both in human evolution and neurodegenerative disease.

## Materials and Methods

### Antibodies

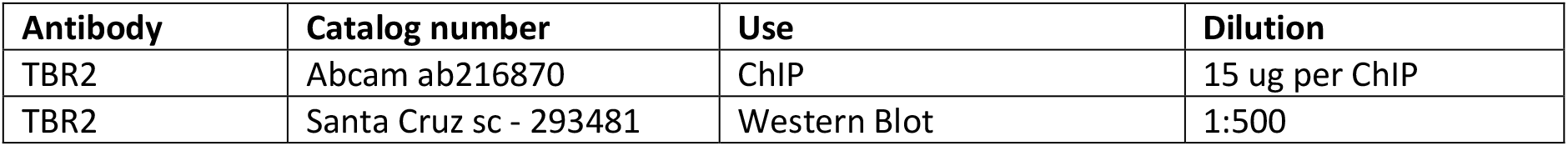
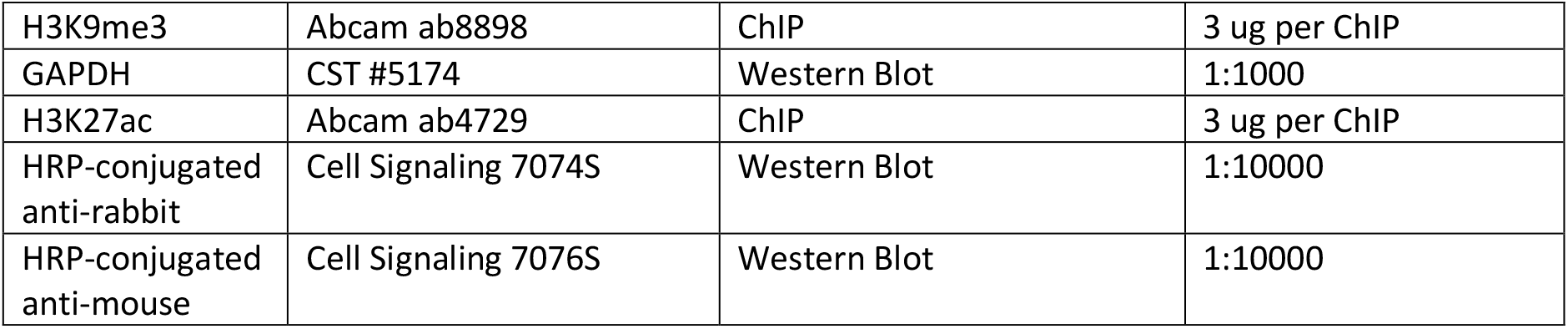

| Antibody | Catalog number | Use | Dilution |
| --- | --- | --- | --- |
| TBR2 | Abcam ab216870 | ChIP | 15 ug per ChIP |
| TBR2 | Santa Cruz sc - 293481 | Western Blot | 1:500 |
| H3K9me3 | Abcam ab8898 | ChIP | 3 ug per ChIP |
| GAPDH | CST #5174 | Western Blot | 1:1000 |
| H3K27ac | Abcam ab4729 | ChIP | 3 ug per ChIP |
| HRP-conjugated anti-rabbit | Cell Signaling 7074S | Western Blot | 1:10000 |
| HRP-conjugated anti-mouse | Cell Signaling 7076S | Western Blot | 1:10000 |

### Human and Chimpanzee iPSC cultures

The human male iPSC line denoted as SV20 was obtained from the University of Pennsylvania, where it was generated, and validated by the expression of pluripotency markers and differentiation into various cell types in multiple studies (Pagliaroli et al., 2021; Pashos et al., 2017; Yang et al., 2015; Zhang et al., 2015). The human female iPSC line GM 23716 was obtained from the Coriell Institute for Medical Research (Camden, NJ) and validated by the expression of pluripotency markers and differentiation into cranial neural crest cells in a previous study (Pagliaroli et al., 2021). The human female iPSC line 21792 and all three of the chimpanzee iPSC lines were obtained from the laboratory of Yoav Gilad at the University of Chicago and validated in previous studies (Gallego Romero et al., 2015; Ward et al., 2018)

The iPSC lines were expanded in feeder-free, serum-free mTeSR™1 medium (STEMCELL Technologies). Cells were passaged ∼1:10 at 80% confluency using ReLeSR (STEMCELL Technologies) and small cell clusters (50–200 cells) were subsequently plated on tissue culture dishes coated overnight with Geltrex™ LDEV-Free hESC-qualified Reduced Growth Factor Basement Membrane Matrix (Fisher-Scientific).

### hpIPC Differentiation

The iPSC lines were differentiated into hpIPCs as previously described (Yu et al. 2014). 3 batches comprised of one human and one chimpanzee iPSC line each were cultured until approximately 50-70% confluence was reached and then treated with the hpIPC media for five days prior to collection for RNA-seq, ATAC-seq, ChIP-seq or immunofluorescence. The hpIPC media consists of DMEM/F12, 0.5X N2, 0.5X B27, DKK-1, cyclopamine, Noggin, and SB431542 as well as the antibiotic Penn-Strep.

### Western Blot

For total lysate, cells were harvested and washed three times in 1X PBS and lysed in RIPA buffer (50mM Tris-HCl pH7.5, 150mM NaCl, 1% Igepal, 0.5% sodium deoxycholate, 0.1% SDS, 500uM DTT) with proteases inhibitors. Twenty μg of whole cell lysate were loaded in Novex WedgeWell 4-20% Tris-Glycine Gel (Invitrogen) and separated through gel electrophoresis (SDS-PAGE) Tris-Glycine-SDS buffer (Invitrogen). The proteins were then transferred to ImmunBlot PVDF membranes (ThermoFisher) for antibody probing. Membranes were incubated with 10% BSA in TBST for 30 minutes at room temperature (RT), then incubated for variable times with the suitable antibodies diluted in 5% BSA in 1X TBST, washed with TBST and incubated with a dilution of 1:10000 of secondary antibody for one hour at RT. The antibody was then visualized using Super Signal West Dura Extended Duration Substrat (ThermoFisher) and imaged with Amersham Imager 680. Used antibodies are listed in the “Antibodies” section of the methods.

### Immunofluorescence

Upon fixation (4% PFA for 10 minutes), cells were first treated with 10 mM sodium citrate (pH = 6) for 10 minutes at 95 C. The cells were then permeabilized in blocking solution (0.1% Triton X-100, re PBS, 5% normal donkey serum) for one hour at room temperature and then incubated overnight at 4 C with the primary antibody of interest. Subsequently, the cells were treated with blocking solution for 30 minutes at room temperature and then fluorophore-conjugated secondary antibodies for 2 hours at room temperature. The cell nuclei were stained with 4,6-diamidine-2-phenylindole dihydrochloride (DAPI; Sigma-Aldrich; 50 mg/ml in PBS for 15 min at RT). Immunostained cells analyzed with confocal microscopy, using a Nikon A1R+. Images were captured with a 40X objectives and a pinhole of 1.2 Airy unit. Analyses were performed in sequential scanning mode to rule out cross-bleeding between channels. Used antibodies are listed in the “Antibodies” section of the methods.

### Real-time quantitative polymerase chain reaction (RT-qPCR)

Cells were lysed in Tri-reagent and RNA was extracted using the Direct-zol RNA MiniPrep kit (Zymo research). 600ng of template RNA was retrotranscribed into cDNA using RevertAid first strand cDNA synthesis kit (Thermo Scientific) according to manufacturer directions. 15ng of cDNA were used for each real-time quantitative PCR reaction with 0.1 μM of each primer, 10 μL of PowerUp™ SYBR™ Green Master Mix (Applied Biosystems) in a final volume of 20 μl, using QuantStudio 3 Real-Time PCR System (Applied Biosystem). Thermal cycling parameters were set as following: 3 minutes at 95°C, followed by 40 cycles of 10 s at 95°C, 20 s at 63°C followed by 30 s at 72°C. Each sample was run in triplicate. 18S rRNA was used as normalizer. Primer sequences are reported in Supplementary Data 1.

### ChIP-Seq and ChiP-qPCR

Samples from different conditions were processed together to prevent batch effects. 15 million cells were cross-linked with 1% formaldehyde for 5 min at room temperature, quenched with 125mM glycine, harvested and washed twice with 1× PBS. The pellet was resuspended in ChIP lysis buffer (150 mM NaCl, 1% Triton X-100, 0,7% SDS, 500 μM DTT, 10 mM Tris-HCl, 5 mM EDTA) and chromatin was sheared to an average length of 200–500 bp, using a Covaris S220 Ultrasonicator. The chromatin lysate was diluted with SDS-free ChIP lysis buffer. For ChIP-seq, 10 µg of antibody (3 µg for H3K27ac) was added to 5 µg of sonicated chromatin along with Dynabeads Protein A magnetic beads (Invitrogen) and incubated at 4 °C overnight. On day 2, beads were washed twice with each of the following buffers: Mixed Micelle Buffer (150 mM NaCl, 1% Triton X-100, 0.2% SDS, 20 mM Tris-HCl, 5 mM EDTA, 65% sucrose), Buffer 500 (500 mM NaCl, 1% Triton X-100, 0.1% Na deoxycholate, 25 mM HEPES, 10 mM Tris-HCl, 1 mM EDTA), LiCl/detergent wash (250 mM LiCl, 0.5% Na deoxycholate, 0.5% NP-40, 10 mM Tris-HCl, 1 mM EDTA) and a final wash was performed with 1× TE. Finally, beads were resuspended in 1× TE containing 1% SDS and incubated at 65 °C for 10 min to elute immunocomplexes. Elution was repeated twice, and the samples were further incubated overnight at 65 °C to reverse cross-linking, along with the untreated input (5% of the starting material). On day 3, after treatment with 0.5 mg/ml Proteinase K for 1h at 65 °C, DNA was purified with Zymo ChIP DNA Clear Concentrator kit and quantified with QUBIT.

For all ChIP-seq experiments, barcoded libraries were made with NEB ULTRA II DNA Library Prep Kit for Illumina, and sequenced on Illumina NextSeq 500, producing 100 bp PE reads.

For ChIP-qPCR, on day 1 the sonicated lysate was aliquot into single immunoprecipitations of 2.5 × 10^6^ cells each. A specific antibody or a total rabbit IgG control was added to the lysate along with Protein A magnetic beads (Invitrogen) and incubated at 4 °C overnight. On day3, ChIP eluates and input were assayed by real-time quantitative PCR in a 20 μl reaction with the following: 0.4 μM of each primer, 10 μl of PowerUp SYBR Green (Applied Biosystems), and 5 μl of template DNA (corresponding to 1/40 of the elution material) using the fast program on QuantStudio qPCR machine (Applied Biosystems). Thermal cycling parameters were: 20sec at 95 °C, followed by 40 cycles of 1sec at 95°C, 20sec at 60°C. Used antibodies are listed in the “Antibodies” section of the methods.

### ChIP-seq Analyses

After removing the adapters with TrimGalore!, the sequences were aligned to the reference hg19, using Burrows Wheeler Alignment tool (BWA), with the MEM algorithm^64^. Uniquely mapping aligned reads were filtered based on mapping quality (MAPQ > 10) to restrict our analysis to higher quality and likely uniquely mapped reads, and PCR duplicates were removed. We called peaks for each individual using MACS2^65^ (H3K27ac) or Homer^66^, at 5% FDR, with default parameters.

### RNA-Seq

Cells were lysed in Tri-reagent (Zymo research) and total RNA was extracted using Quick-RNA Miniprep kit (Zymo research) according to the manufacturer’s instructions. RNA was further quantified using DeNovix DS-11 Spectrophotometer while the RNA integrity was checked on Bioanalyzer 2100 (Agilent). Only samples with RIN value above 8.0 were used for transcriptome analysis. RNA libraries were prepared using 1 μg of total RNA input using NEBNext® Poly(A) mRNA Magnetic Isolation Module, NEBNext® UltraTM II Directional RNA Library Prep Kit for Illumina® and NEBNext® UltraTM II DNA Library Prep Kit for Illumina® according to the manufacturer’s instructions (New England Biolabs).

### Single-cell RNA-Seq

Cells from both species at each time were first incubated with Accutase at 37 C for 7 minutes. The cells were collected with DMEM/F12 and centrifuged for 5 minutes at 150 xg. The cells were resuspended in 10% FBS in DMEM and strained with a 40 um cell strainer to create a single-cell suspension. After confirming >90% viability with a Countess III, the cells were processed with the 10X Genomics Cell Multiplexing Oligo protocol.

### Single-cell RNA-Seq analyses

10X Cell Ranger (Zheng et al., 2017) was used to demultiplex and map the scRNA-seq data, with the tools cellranger multi and cellranger mkfastq. Seurat 4 (Hao et al., 2021) was used for individual analysis of scRNA-seq data, as well as integration of human and chimpanzee datasets. The human genes used to classify the cells into the G1/S/G2M phases of the cell cycle were obtained from https://github.com/hbctraining/scRNA-seq/blob/master/lessons/06_SC_SCT_and_integration.md

### RNA-Seq Analyses

After removing the adapters with TrimGalore!, Kallisto (Bray et al., 2016) was used to count reads mapping to each gene. We analyzed differential gene expression levels with DESeq2 (Love et al., 2014), with the following model: design = ∼condition, where condition indicates either Human or Chimpanzee.

### ATAC-Seq

For ATAC-Seq experiments, 50,000 cells per condition were processed as described in the original ATAC-seq protocol paper (Buenrostro et al. 2013). ATAC-seq data were processed with the same pipeline described for ChIP-seq, with one modification: all mapped reads were offset by +4 bp for the forward-strand and -5 bp for the reverse-strand. Peaks were called using MACS2 (Zhang et al., 2008, 2).

### Generation of the NCCIT-dCas9KRAB-SVAsgRNA Stable Cell Line

This cell line was generated in our recent study (Barnada et al., 2022). Briefly, a dCas9-KRAB was cloned into a piggyBac transposon containing ampicillin and puromycin resistance (Addgene) which was obtained from the Wysocka Lab at Stanford University. The piggyBac dCas9-KRAB doxycycline-inducible plasmid, along with a piggyBac transposase (Cell Signaling), were transfected into NCCIT cells (ATCC) at ∼70% confluency using a 6:1 ratio of Fugene HD (Promega) for 48 hours in ATCC-formulated RPMI media. Two days post-transfection, the media was changed and the transfected cells were selected for using 1µg puromycin per 1mL media. A piggyBac transposon plasmid containing two sgRNAs (SVAsgRNA1: 5’CTCCCTAATCTCAAGTACCC 3’ and SVAsgRNA2: 5’ TGTTTCAGAGAGCACGGGGT 3’; Integrated DNA Technologies) targeting ∼80% of all annotated SVAs in humans (Pontis et al. 2019), along with a piggyBac transposase, were transfected into the NCCIT-dCas9KRAB cells using a 6:1 ratio of Fugene HD for 48 hours in ATCC-formulated RPMI media. Two days post-transfection, the media was changed and the transfected cells were selected for using 400µg geneticin per 1mL of media in addition to 1µg puromycin per 1mL media. The NCCIT-dCas9KRAB-SVAsgRNA cell line was maintained in ATCC-formulated RPMI media supplemented with 10% tet-free FBS, 1% L-glutamine, 1µg/mL puromycin, and 400µg/mL geneticin and incubated at 5% CO_2_, 20% O_2_ at 37°C.

### Retinoic Acid-induced neuronal differentiation and CRISPR-interference of NCCIT-dCas9KRAB-SVAsgRNA cells

The NCCIT-dCas9KRAB-SVAsgRNA cells, at ∼20% confluency, were treated with 10µM retinoic acid (RA) per 10mL media for 1 week to induce neuronal differentiation. At day 4, the media was refreshed and the cells were additionally treated with 2µg doxycycline per 1mL of media for 3 days. The cells were collected on day 7 of RA treatment (day 3 of doxycycline treatment) for qPCR and genomic experiments. Expression of the doxycycline-inducible dCas9 was verified via western blot.

### Statistical and genomic analyses

All statistical analyses were performed using R v3.3.1 or Graphpad Prism version 9.2.0 for Mac OS X. BEDtools v2.27.1 (Quinlan and Hall, 2010) was used for genomic analyses. Pathway analysis was performed with Ingenuity Pathway Analysis Suite (QIAGEN Inc., https://www.qiagenbioinformatics.com/products/ingenuity-pathway-analysis). Motif analyses were performed using the MEME-Suite (Bailey et al., 2015), and specifically with the Meme-ChIP application. Fasta files of the regions of interest were produced using BEDTools v2.27.1. Shuffled input sequences were used as background. E-values < 0.001 were used as threshold for significance. Orthologous ATAC-seq regions were identified using the University of California Santa Cruz Genome Browser tool LiftOver.

## Acknowledgements

The authors are grateful to Dr. Yoav Gilad (University of Chicago) and with the Yerkes National Primate Research Center (Emory University, Atlanta) for providing all the chimpanzee lines and one of the human lines, and to Dr. Joanna Wysocka’s lab, and in particular Dr. Raquel Fueyo (Stanford University), for providing CRISPR reagents and critical advice. The authors thank the Thomas Jefferson Stem Cell and Regenerative Neuroscience Center for the support in the process of optimization of the iPSC differentiation. The authors are grateful to Trizzino lab members Luca Pagliaroli, PhD, Chiara Scopa, PhD and Connor Ott for help with specific analyses and experiments. The authors thank the Genomic Facility at The Wistar Institute (Philadelphia, PA) for the Next Generation Illumina Sequencing. The PAX6 antibody developed by Kawakami, A. was obtained from the Developmental Hybridoma Bank, created by the NICHD of the NIH and maintained at The University of Iowa, Department of Biology, Iowa City, IA 52242.

## Data availability

The original genome-wide data generated in this study have been deposited in the GEO database under accession code GSE189347.

## Author contributions

MT and SP designed the project. SP performed most of the experiments. SMB carried out the CRISPR experiments in NCCIT cells. SP and MT analyzed the data and wrote the manuscript. CL and JIM supervised SP in the scRNA-seq analyses. All the authors read and approved the manuscript.

## Figure legends

**Supplemental Figure 1.**
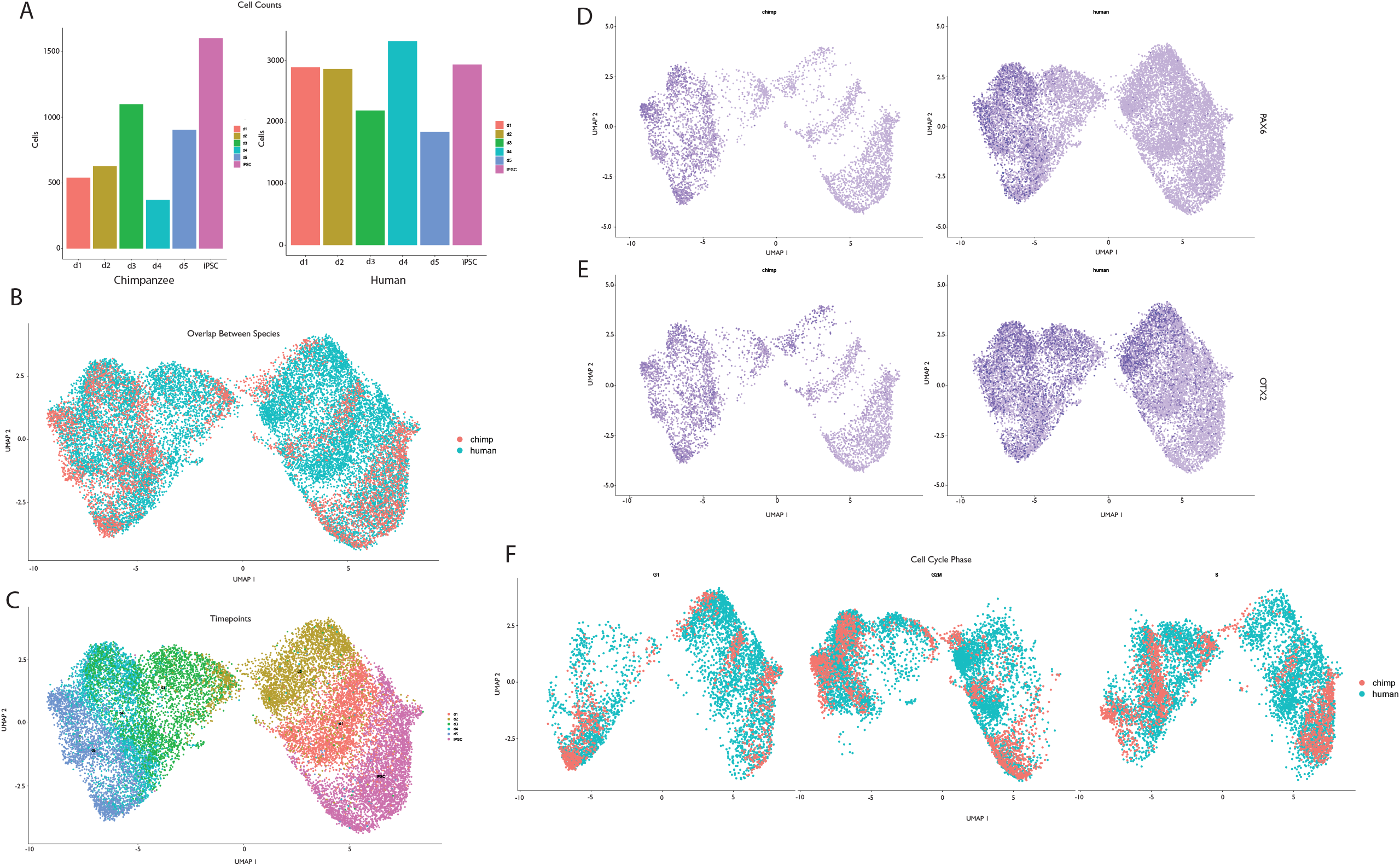
Single Cell RNA-seq Analysis of Human and Chimpanzee hpIPC Differentiation. (A) Number of cells assayed at each time point in each species (B) Overlap of all human and chimpanzee cells in a single plot. Chimpanze cells are depicted in red and human cells are depicted in blue. (C) Overlap of all human and chimpanzee cells across all six time points in a single plot. (D) Expression of PAX6 across all six time points, human cells are depicted on the right and chimpanzee cells are depicted on the left. (E) Expression of OTX2 across all six time points, human cells are depicted on the left and chimpanzee cells are depicted on the right (F) Overlap of human and chimpanzee cells at G1, G2M and S phases of the cell cycle.

## References

Abrajano, J. J., Qureshi, I. A., Gokhan, S., Zheng, D., Bergman, A. and Mehler, M. F. (2009). REST and CoREST Modulate Neuronal Subtype Specification, Maturation and Maintenance. PLOS ONE 4, e7936.

Agoglia, R. M., Sun, D., Birey, F., Yoon, S.-J., Miura, Y., Sabatini, K., Pașca, S. P. and Fraser, H. B. (2021). Primate cell fusion disentangles gene regulatory divergence in neurodevelopment. Nature 592, 421–427.

Andersen, J., Urbán, N., Achimastou, A., Ito, A., Simic, M., Ullom, K., Martynoga, B., Lebel, M., Göritz, C., Frisén, J., et al. (2014). A Transcriptional Mechanism Integrating Inputs from Extracellular Signals to Activate Hippocampal Stem Cells. Neuron 83, 1085–1097.

Arnold, S. J., Huang, G.-J., Cheung, A. F. P., Era, T., Nishikawa, S.-I., Bikoff, E. K., Molnar, Z., Robertson, E. J. and Groszer, M. (2008). The T-box transcription factor Eomes/Tbr2 regulates neurogenesis in the cortical subventricular zone. Genes & Development 22, 2479–2484.

Aruga, J. (2004). The role of Zic genes in neural development. Mol Cell Neurosci 26, 205–221.

Averaimo, S., Gritti, M., Barini, E., Gasparini, L. and Mazzanti, M. (2014). CLIC1 functional expression is required for cAMP-induced neurite elongation in post-natal mouse retinal ganglion cells. Journal of Neurochemistry 131, 444–456.

Bailey, T. L., Johnson, J., Grant, C. E. and Noble, W. S. (2015). The MEME Suite. Nucleic Acids Research 43, W39–W49.

Barger, N., Hanson, K. L., Teffer, K., Schenker-Ahmed, N. M. and Semendeferi, K. (2014). Evidence for evolutionary specialization in human limbic structures. Front. Hum. Neurosci. 8,.

Barnada, S. M., Isopi, A., Tejada-Martinez, D., Goubert, C., Patoori, S., Pagliaroli, L., Tracewell, M. and Trizzino, M. (2022). Genomic features underlie the co-option of SVA transposons as cis-regulatory elements in human pluripotent stem cells. PLOS Genetics 18, e1010225.

Bekinschtein, P., Katche, C., Slipczuk, L. N., Igaz, L. M., Cammarota, M., Izquierdo, I. and Medina, J. H. (2007). mTOR signaling in the hippocampus is necessary for memory formation. Neurobiology of Learning and Memory 87, 303–307.

Bray, N. L., Pimentel, H., Melsted, P. and Pachter, L. (2016). Near-optimal probabilistic RNA-seq quantification. Nat Biotechnol 34, 525–527.

Bulfone, A., Martinez, S., Marigo, V., Campanella, M., Basile, A., Quaderi, N., Gattuso, C., Rubenstein, J. L. R. and Ballabio, A. (1999). Expression pattern of the Tbr2 (Eomesodermin) gene during mouse and chick brain development. Mechanisms of Development 84, 133–138.

Burgess, N., Maguire, E. A. and O’Keefe, J. (2002). The Human Hippocampus and Spatial and Episodic Memory. Neuron 35, 625–641.

Chen, X., Wang, S., Zhou, Y., Han, Y., Li, S., Xu, Q., Xu, L., Zhu, Z., Deng, Y., Yu, L., et al. (2018). Phf8 histone demethylase deficiency causes cognitive impairments through the mTOR pathway. Nat Commun 9, 114.

Cheval, H., Chagneau, C., Levasseur, G., Veyrac, A., Faucon-Biguet, N., Laroche, S. and Davis, S. (2012). Distinctive features of Egr transcription factor regulation and DNA binding activity in CA1 of the hippocampus in synaptic plasticity and consolidation and reconsolidation of fear memory. Hippocampus 22, 631–642.

Chuong, E. B., Rumi, M. A. K., Soares, M. J. and Baker, J. C. (2013). Endogenous retroviruses function as species-specific enhancer elements in the placenta. Nat Genet 45, 325–329.

Chuong, E. B., Elde, N. C. and Feschotte, C. (2016). Regulatory evolution of innate immunity through co-option of endogenous retroviruses. Science 351, 1083–1087.

Cipriani, S., Nardelli, J., Verney, C., Delezoide, A.-L., Guimiot, F., Gressens, P. and Adle-Biassette, H. (2016). Dynamic Expression Patterns of Progenitor and Pyramidal Neuron Layer Markers in the Developing Human Hippocampus. Cereb. Cortex 26, 1255–1271.

Coluccio, A., Ecco, G., Duc, J., Offner, S., Turelli, P. and Trono, D. (2018). Individual retrotransposon integrants are differentially controlled by KZFP/KAP1-dependent histone methylation, DNA methylation and TET-mediated hydroxymethylation in naïve embryonic stem cells. Epigenetics & Chromatin 11, 7.

Cosby, R. L., Judd, J., Zhang, R., Zhong, A., Garry, N., Pritham, E. J. and Feschotte, C. (2021). Recurrent evolution of vertebrate transcription factors by transposase capture. Science 371, eabc6405.

Coufal, N. G., Garcia-Perez, J. L., Peng, G. E., Yeo, G. W., Mu, Y., Lovci, M. T., Morell, M., O’Shea, K. S., Moran, J. V. and Gage, F. H. (2009). L1 retrotransposition in human neural progenitor cells. Nature 460, 1127–1131.

Crawley, J. N., Heyer, W.-D. and LaSalle, J. M. (2016). Autism and Cancer Share Risk Genes, Pathways, and Drug Targets. Trends in Genetics 32, 139–146.

Datson, N. a., Morsink, M. c., Steenbergen, P. j., Aubert, Y., Schlumbohm, C., Fuchs, E. and de Kloet, E. r. (2009). A molecular blueprint of gene expression in hippocampal subregions CA1, CA3, and DG is conserved in the brain of the common marmoset. Hippocampus 19, 739–752.

Davis, L., Onn, I. and Elliott, E. (2018). The emerging roles for the chromatin structure regulators CTCF and cohesin in neurodevelopment and behavior. Cell. Mol. Life Sci. 75, 1205–1214.

del Rosario, R. C. H., Rayan, N. A. and Prabhakar, S. (2014). Noncoding origins of anthropoid traits and a new null model of transposon functionalization. Genome Res 24, 1469–1484.

Du, J., Leung, A., Trac, C., Lee, M., Parks, B. W., Lusis, A. J., Natarajan, R. and Schones, D. E. (2016). Chromatin variation associated with liver metabolism is mediated by transposable elements. Epigenetics & Chromatin 9, 28.

Duyckaerts, C., Delatour, B. and Potier, M.-C. (2009). Classification and basic pathology of Alzheimer disease. Acta Neuropathol 118, 5–36.

Edler, M. K., Sherwood, C. C., Meindl, R. S., Hopkins, W. D., Ely, J. J., Erwin, J. M., Mufson, E. J., Hof, P. R. and Raghanti, M. A. (2017). Aged chimpanzees exhibit pathologic hallmarks of Alzheimer’s disease. Neurobiology of Aging 59, 107–120.

Eichenbaum, H. (2017a). The role of the hippocampus in navigation is memory. Journal of Neurophysiology 117, 1785–1796.

Eichenbaum, H. (2017b). Prefrontal–hippocampal interactions in episodic memory. Nat Rev Neurosci 18, 547–558.

Emera, D., Yin, J., Reilly, S. K., Gockley, J. and Noonan, J. P. (2016). Origin and evolution of developmental enhancers in the mammalian neocortex. PNAS 113, E2617–E2626.

Enard, W. (2011). FOXP2 and the role of cortico-basal ganglia circuits in speech and language evolution. Current Opinion in Neurobiology 21, 415–424.

Enard, W., Khaitovich, P., Klose, J., Zöllner, S., Heissig, F., Giavalisco, P., Nieselt-Struwe, K., Muchmore, E., Varki, A., Ravid, R., et al. (2002). Intra- and Interspecific Variation in Primate Gene Expression Patterns. Science 296, 340–343.

Englund, C. (2005). Pax6, Tbr2, and Tbr1 Are Expressed Sequentially by Radial Glia, Intermediate Progenitor Cells, and Postmitotic Neurons in Developing Neocortex. Journal of Neuroscience 25, 247–251.

Ferrón, S. R., Charalambous, M., Radford, E., McEwen, K., Wildner, H., Hind, E., Morante-Redolat, J. M., Laborda, J., Guillemot, F., Bauer, S. R., et al. (2011). Postnatal loss of Dlk1 imprinting in stem cells and niche astrocytes regulates neurogenesis. Nature 475, 381–385.

Finch, C. E. and Austad, S. N. (2015). Commentary: is Alzheimer’s disease uniquely human? Neurobiology of Aging 36, 553–555.

Florio, M. and Huttner, W. B. (2014). Neural progenitors, neurogenesis and the evolution of the neocortex. Development 141, 2182–2194.

Fortress, A. M., Fan, L., Orr, P. T., Zhao, Z. and Frick, K. M. (2013). Estradiol-induced object recognition memory consolidation is dependent on activation of mTOR signaling in the dorsal hippocampus. Learn. Mem. 20, 147–155.

Fuentes, D. R., Swigut, T. and Wysocka, J. (2018). Systematic perturbation of retroviral LTRs reveals widespread long-range effects on human gene regulation. eLife 7, e35989.

Gallego Romero, I., Pavlovic, B. J., Hernando-Herraez, I., Zhou, X., Ward, M. C., Banovich, N. E., Kagan, C. L., Burnett, J. E., Huang, C. H., Mitrano, A., et al. (2015). A panel of induced pluripotent stem cells from chimpanzees: a resource for comparative functional genomics. eLife 4, e07103.

Gokhman, D., Agoglia, R. M., Kinnebrew, M., Gordon, W., Sun, D., Bajpai, V. K., Naqvi, S., Chen, C., Chan, A., Chen, C., et al. (2021). Human–chimpanzee fused cells reveal cis -regulatory divergence underlying skeletal evolution. Nature Genetics 53, 467–476.

Graber, T. E., McCamphill, P. K. and Sossin, W. S. (2013). A recollection of mTOR signaling in learning and memory. Learn. Mem. 20, 518–530.

Hao, Y., Hao, S., Andersen-Nissen, E., Mauck, W. M., Zheng, S., Butler, A., Lee, M. J., Wilk, A. J., Darby, C., Zager, M., et al. (2021). Integrated analysis of multimodal single-cell data. Cell 184, 3573-3587.e29.

Hevner, R. F. (2019). Intermediate progenitors and Tbr2 in cortical development. J. Anat. 235, 616–625.

Hickey, S. L., Berto, S. and Konopka, G. (2019). Chromatin Decondensation by FOXP2 Promotes Human Neuron Maturation and Expression of Neurodevelopmental Disease Genes. Cell Reports 27, 1699-1711.e9.

Hodge, R. D., Nelson, B. R., Kahoud, R. J., Yang, R., Mussar, K. E., Reiner, S. L. and Hevner, R. F. (2012). Tbr2 Is Essential for Hippocampal Lineage Progression from Neural Stem Cells to Intermediate Progenitors and Neurons. Journal of Neuroscience 32, 6275–6287.

Ito, J., Kimura, I., Soper, A., Coudray, A., Koyanagi, Y., Nakaoka, H., Inoue, I., Turelli, P., Trono, D. and Sato, K. Endogenous retroviruses drive KRAB zinc-finger protein family expression for tumor suppression. Science Advances 6, eabc3020.

Ivancevic, A. and Chuong, E. B. (2020). Transposable elements teach T cells new tricks. PNAS 117, 9145– 9147.

Jacques, P.-É., Jeyakani, J. and Bourque, G. (2013). The Majority of Primate-Specific Regulatory Sequences Are Derived from Transposable Elements. PLOS Genetics 9, e1003504.

Jönsson, M. E., Ludvik Brattås, P., Gustafsson, C., Petri, R., Yudovich, D., Pircs, K., Verschuere, S., Madsen, S., Hansson, J., Larsson, J., et al. (2019). Activation of neuronal genes via LINE-1 elements upon global DNA demethylation in human neural progenitors. Nat Commun 10, 3182.

Judd, J., Sanderson, H. and Feschotte, C. (2021). Evolution of mouse circadian enhancers from transposable elements. Genome Biol 22, 193.

Kamboh, M. I., Fan, K.-H., Yan, Q., Beer, J. C., Snitz, B. E., Wang, X., Chang, C.-C. H., Demirci, F. Y., Feingold, E. and Ganguli, M. (2019). Population-based genome-wide association study of cognitive decline in older adults free of dementia: identification of a novel locus for the attention domain. Neurobiol Aging 84, 239.e15-239.e24.

Kimura, N., Nakashima, K., Ueno, M., Kiyama, H. and Taga, T. (1999). A novel mammalian T-box-containing gene, Tbr2, expressed in mouse developing brain. Developmental Brain Research 115, 183–193.

King, M. and Wilson, A. (1975). Evolution at two levels in humans and chimpanzees. Science 188, 107– 116.

Kittappa, R., Chang, W. W., Awatramani, R. B. and McKay, R. D. G. (2007). The foxa2 Gene Controls the Birth and Spontaneous Degeneration of Dopamine Neurons in Old Age. PLOS Biology 5, e325.

Kohane, A. A. and Wood, T. R. (2021). Neurodevelopmental clustering of gene expression identifies lipid metabolism genes associated with neuroprotection and neurodegeneration.

Kolehmainen, J., Black, G. C. M., Saarinen, A., Chandler, K., Clayton-Smith, J., Träskelin, A.-L., Perveen, R., Kivitie-Kallio, S., Norio, R., Warburg, M., et al. (2003). Cohen syndrome is caused by mutations in a novel gene, COH1, encoding a transmembrane protein with a presumed role in vesicle-mediated sorting and intracellular protein transport. Am J Hum Genet 72, 1359–1369.

Liégeois, F. J., Hildebrand, M. S., Bonthrone, A., Turner, S. J., Scheffer, I. E., Bahlo, M., Connelly, A. and Morgan, A. T. (2016). Early neuroimaging markers of FOXP2 intragenic deletion. Sci Rep 6, 35192.

Love, M. I., Huber, W. and Anders, S. (2014). Moderated estimation of fold change and dispersion for RNA-seq data with DESeq2. Genome Biology 15, 550.

Lynch, V. J., Leclerc, R. D., May, G. and Wagner, G. P. (2011). Transposon-mediated rewiring of gene regulatory networks contributed to the evolution of pregnancy in mammals. Nat Genet 43, 1154–1159.

Lynch, V. J., Nnamani, M. C., Kapusta, A., Brayer, K., Plaza, S. L., Mazur, E. C., Emera, D., Sheikh, S. Z., Grützner, F., Bauersachs, S., et al. (2015). Ancient Transposable Elements Transformed the Uterine Regulatory Landscape and Transcriptome during the Evolution of Mammalian Pregnancy. Cell Rep 10, 551–561.

MacDermot, K. D., Bonora, E., Sykes, N., Coupe, A.-M., Lai, C. S. L., Vernes, S. C., Vargha-Khadem, F., McKenzie, F., Smith, R. L., Monaco, A. P., et al. (2005). Identification of FOXP2 Truncation as a Novel Cause of Developmental Speech and Language Deficits. The American Journal of Human Genetics 76, 1074–1080.

Mandal, C., Jung, K. H., Kang, S. C., Choi, M. R., Park, K. S., Chung, I. Y. and Chai, Y. G. (2015). Knocking down of UTX in NCCIT cells enhance cell attachment and promote early neuronal cell differentiation. BioChip J 9, 182–193.

Marchetto, M. C., Hrvoj-Mihic, B., Kerman, B. E., Yu, D. X., Vadodaria, K. C., Linker, S. B., Narvaiza, I., Santos, R., Denli, A. M., Mendes, A. P., et al. (2019). Species-specific maturation profiles of human, chimpanzee and bonobo neural cells. eLife 8, e37527.

Martínez-Cerdeño, V., Noctor, S. C. and Kriegstein, A. R. (2006). The role of intermediate progenitor cells in the evolutionary expansion of the cerebral cortex. Cereb Cortex 16 Suppl 1, i152–161.

Mathys, H., Davila-Velderrain, J., Peng, Z., Gao, F., Mohammadi, S., Young, J. Z., Menon, M., He, L., Abdurrob, F., Jiang, X., et al. (2019). Single-cell transcriptomic analysis of Alzheimer’s disease. Nature 570, 332–337.

Miao, B., Fu, S., Lyu, C., Gontarz, P., Wang, T. and Zhang, B. (2020). Tissue-specific usage of transposable element-derived promoters in mouse development. Genome Biol 21, 255.

Mihalas, A. B., Elsen, G. E., Bedogni, F., Daza, R. A. M., Ramos-Laguna, K. A., Arnold, S. J. and Hevner, R. F. (2016). Intermediate Progenitor Cohorts Differentially Generate Cortical Layers and Require Tbr2 for Timely Acquisition of Neuronal Subtype Identity. Cell Reports 16, 92–105.

Mika, K., Marinić, M., Singh, M., Muter, J., Brosens, J. J. and Lynch, V. J. (2021). Evolutionary transcriptomics implicates new genes and pathways in human pregnancy and adverse pregnancy outcomes. eLife 10, e69584.

Modzelewski, A. J., Shao, W., Chen, J., Lee, A., Qi, X., Noon, M., Tjokro, K., Sales, G., Biton, A., Anand, A., et al. (2021). A mouse-specific retrotransposon drives a conserved Cdk2ap1 isoform essential for development. Cell 184, 5541-5558.e22.

Mora-Bermúdez, F., Badsha, F., Kanton, S., Camp, J. G., Vernot, B., Köhler, K., Voigt, B., Okita, K., Maricic, T., He, Z., et al. (2016). Differences and similarities between human and chimpanzee neural progenitors during cerebral cortex development. eLife 5, e18683.

Mukherjee, D., Gonzales, B. J., Ashwal-Fluss, R., Turm, H., Groysman, M. and Citri, A. (2021). Egr2 induction in spiny projection neurons of the ventrolateral striatum contributes to cocaine place preference in mice. eLife 10, e65228.

Okhovat, M., Nevonen, K. A., Davis, B. A., Michener, P., Ward, S., Milhaven, M., Harshman, L., Sohota, A., Fernandes, J. D., Salama, S. R., et al. (2020). Co-option of the lineage-specific LAVA retrotransposon in the gibbon genome. PNAS 117, 19328–19338.

Pagliaroli, L., Porazzi, P., Curtis, A. T., Scopa, C., Mikkers, H. M. M., Freund, C., Daxinger, L., Deliard, S., Welsh, S. A., Offley, S., et al. (2021). Inability to switch from ARID1A-BAF to ARID1B-BAF impairs exit from pluripotency and commitment towards neural crest formation in ARID1B-related neurodevelopmental disorders. Nat Commun 12, 6469.

Pashos, E. E., Park, Y., Wang, X., Raghavan, A., Yang, W., Abbey, D., Peters, D. T., Arbelaez, J., Hernandez, M., Kuperwasser, N., et al. (2017). Large, Diverse Population Cohorts of hiPSCs and Derived Hepatocyte-like Cells Reveal Functional Genetic Variation at Blood Lipid-Associated Loci. Cell Stem Cell 20, 558-570.e10.

Playfoot, C. J., Duc, J., Sheppard, S., Dind, S., Coudray, A., Planet, E. and Trono, D. (2021). Transposable elements and their KZFP controllers are drivers of transcriptional innovation in the developing human brain. Genome Res. 31, 1531–1545.

Poirier, R., Cheval, H., Mailhes, C., Charnay, P., Davis, S. and Laroche, S. (2007). Paradoxical Role of an Egr Transcription Factor Family Member, Egr2/Krox20, in Learning and Memory. Front Behav Neurosci 1, 6.

Pontious, A., Kowalczyk, T., Englund, C. and Hevner, R. F. (2008). Role of intermediate progenitor cells in cerebral cortex development. Dev Neurosci 30, 24–32.

Pontis, J., Planet, E., Offner, S., Turelli, P., Duc, J., Coudray, A., Theunissen, T. W., Jaenisch, R. and Trono, D. (2019). Hominoid-Specific Transposable Elements and KZFPs Facilitate Human Embryonic Genome Activation and Control Transcription in Naive Human ESCs. Cell Stem Cell 24, 724-735.e5.

Quinlan, A. R. and Hall, I. M. (2010). BEDTools: a flexible suite of utilities for comparing genomic features. Bioinformatics 26, 841–842.

Quinn, J. P. and Bubb, V. J. (2014). SVA retrotransposons as modulators of gene expression. Mobile Genetic Elements 4, e32102.

Raivich, G. (2008). c-Jun expression, activation and function in neural cell death, inflammation and repair. J Neurochem 107, 898–906.

Raivich, G. and Behrens, A. (2006). Role of the AP-1 transcription factor c-Jun in developing, adult and injured brain. Prog Neurobiol 78, 347–363.

Rayan, N. A., Del Rosario, R. C. H. and Prabhakar, S. (2016). Massive contribution of transposable elements to mammalian regulatory sequences. Semin Cell Dev Biol 57, 51–56.

Reilly, S. K., Yin, J., Ayoub, A. E., Emera, D., Leng, J., Cotney, J., Sarro, R., Rakic, P. and Noonan, J. P. (2015). Evolutionary genomics. Evolutionary changes in promoter and enhancer activity during human corticogenesis. Science 347, 1155–1159.

Rétaux, S., Bourrat, F., Joly, J.-S. and Hinaux, H. (2013). Perspectives in Evo-Devo of the Vertebrate Brain. In Advances in Evolutionary Developmental Biology, pp. 151–172. John Wiley & Sons, Ltd.

Reuter, M. S., Krumbiegel, M., Schlüter, G., Ekici, A. B., Reis, A. and Zweier, C. (2017). Haploinsufficiency of NR4A2 is associated with a neurodevelopmental phenotype with prominent language impairment. Am J Med Genet A 173, 2231–2234.

Sakamoto, K., Karelina, K. and Obrietan, K. (2011). CREB: a multifaceted regulator of neuronal plasticity and protection. Journal of Neurochemistry 116, 1–9.

Sarkar, A., Mei, A., Paquola, A. C. M., Stern, S., Bardy, C., Klug, J. R., Kim, S., Neshat, N., Kim, H. J., Ku, M., et al. (2018). Efficient Generation of CA3 Neurons from Human Pluripotent Stem Cells Enables Modeling of Hippocampal Connectivity In Vitro. Cell Stem Cell 22, 684-697.e9.

Schmidt, D., Schwalie, P. C., Wilson, M. D., Ballester, B., Gonçalves, A., Kutter, C., Brown, G. D., Marshall, A., Flicek, P. and Odom, D. T. (2012). Waves of retrotransposon expansion remodel genome organization and CTCF binding in multiple mammalian lineages. Cell 148, 335–348.

Sessa, A., Mao, C.-A., Hadjantonakis, A.-K., Klein, W. H. and Broccoli, V. (2008). Tbr2 directs conversion of radial glia into basal precursors and guides neuronal amplification by indirect neurogenesis in the developing neocortex. Neuron 60, 56–69.

Shah, A. H., Gilbert, M., Ivan, M. E., Komotar, R. J., Heiss, J. and Nath, A. (2021). The role of human endogenous retroviruses in gliomas: from etiological perspectives and therapeutic implications. Neuro-Oncology 23, 1647–1655.

Sin, C., Li, H. and Crawford, D. A. (2015). Transcriptional Regulation by FOXP1, FOXP2, and FOXP4 Dimerization. Journal of Molecular Neuroscience 55, 437–448.

Sousa, A. M. M., Zhu, Y., Raghanti, M. A., Kitchen, R. R., Onorati, M., Tebbenkamp, A. T. N., Stutz, B., Meyer, K. A., Li, M., Kawasawa, Y. I., et al. (2017). Molecular and cellular reorganization of neural circuits in the human lineage. Science 358, 1027–1032.

Squire, L. R. (1992). Memory and the Hippocampus: A Synthesis From Findings With Rats, Monkeys, and Humans.

Sundaram, V. and Wysocka, J. (2020). Transposable elements as a potent source of diverse cis-regulatory sequences in mammalian genomes. Philosophical Transactions of the Royal Society B: Biological Sciences 375, 20190347.

Sundaram, V., Cheng, Y., Ma, Z., Li, D., Xing, X., Edge, P., Snyder, M. P. and Wang, T. (2014). Widespread contribution of transposable elements to the innovation of gene regulatory networks. Genome Res 24, 1963–1976.

Sur, D., Kustwar, R. K., Budania, S., Mahadevan, A., Hancks, D. C., Yadav, V., Shankar, S. K. and Mandal, P. K. (2017). Detection of the LINE-1 retrotransposon RNA-binding protein ORF1p in different anatomical regions of the human brain. Mobile DNA 8, 17.

Taher, N., McKenzie, C., Garrett, R., Baker, M., Fox, N. and Isaacs, G. D. (2014). Amyloid-β Alters the DNA Methylation Status of Cell-fate Genes in an Alzheimer’s Disease Model. Journal of Alzheimer’s Disease 38, 831–844.

Thomas, C. A., Paquola, A. C. M. and Muotri, A. R. (2012). LINE-1 Retrotransposition in the Nervous System. Annual Review of Cell and Developmental Biology 28, 555–573.

Tokuyama, M., Kong, Y., Song, E., Jayewickreme, T., Kang, I. and Iwasaki, A. (2018). ERVmap analysis reveals genome-wide transcription of human endogenous retroviruses. PNAS 115, 12565– 12572.

Tomasello, M. and Herrmann, E. (2010). Ape and Human Cognition: What’s the Difference? Current Directions in Psychological Science 19, 3–8.

Trizzino, M., Park, Y., Holsbach-Beltrame, M., Aracena, K., Mika, K., Caliskan, M., Perry, G. H., Lynch, V. J. and Brown, C. D. (2017). Transposable elements are the primary source of novelty in primate gene regulation. Genome Res.

Trizzino, M., Kapusta, A. and Brown, C. D. (2018). Transposable elements generate regulatory novelty in a tissue-specific fashion. BMC Genomics 19, 468.

Turelli, P., Playfoot, C., Grun, D., Raclot, C., Pontis, J., Coudray, A., Thorball, C., Duc, J., Pankevich, E. V., Deplancke, B., et al. (2020). Primate-restricted KRAB zinc finger proteins and target retrotransposons control gene expression in human neurons. Science Advances 6, eaba3200.

Walker, L. C. and Jucker, M. (2017). The Exceptional Vulnerability of Humans to Alzheimer’s Disease. Trends in Molecular Medicine 23, 534–545.

Wang, H., Xing, J., Grover, D., Hedges, D. J., Han, K., Walker, J. A. and Batzer, M. A. (2005). SVA Elements: A Hominid-specific Retroposon Family. Journal of Molecular Biology 354, 994–1007.

Ward, M. C., Zhao, S., Luo, K., Pavlovic, B. J., Karimi, M. M., Stephens, M. and Gilad, Y. (2018). Silencing of transposable elements may not be a major driver of regulatory evolution in primate iPSCs. eLife 7, e33084.

Wei, Y. (2020). Comparative transcriptome analysis of the hippocampus from sleep-deprived and Alzheimer’s disease mice. Genet Mol Biol 43, e20190052.

Wray, G. A. (2007). The evolutionary significance of cis-regulatory mutations. Nat Rev Genet 8, 206–216.

Xie, M., Hong, C., Zhang, B., Lowdon, R. F., Xing, X., Li, D., Zhou, X., Lee, H. J., Maire, C. L., Ligon, K. L., et al. (2013). DNA hypomethylation within specific transposable element families associates with tissue-specific enhancer landscape. Nat Genet 45, 836–841.

Yang, W., Liu, Y., Slovik, K. J., Wu, J. C., Duncan, S. A., Rader, D. J. and Morrisey, E. E. (2015). Generation of iPSCs as a Pooled Culture Using Magnetic Activated Cell Sorting of Newly Reprogrammed Cells. PLoS One 10, e0134995.

Yi, R., Chen, B., Zhao, J., Zhan, X., Zhang, L., Liu, X. and Dong, Q. (2014). Krüppel-like Factor 8 Ameliorates Alzheimer’s Disease by Activating β-Catenin. J Mol Neurosci 52, 231–241.

Yu, D. X., Di Giorgio, F. P., Yao, J., Marchetto, M. C., Brennand, K., Wright, R., Mei, A., Mchenry, L., Lisuk, D., Grasmick, J. M., et al. (2014). Modeling Hippocampal Neurogenesis Using Human Pluripotent Stem Cells. Stem Cell Reports 2, 295–310.

Zhang, Y., Liu, T., Meyer, C. A., Eeckhoute, J., Johnson, D. S., Bernstein, B. E., Nusbaum, C., Myers, R. M., Brown, M., Li, W., et al. (2008). Model-based Analysis of ChIP-Seq (MACS). Genome Biology 9, R137.

Zhang, J., Li, H., Yabut, O., Fitzpatrick, H., D’Arcangelo, G. and Herrup, K. (2010). Cdk5 Suppresses the Neuronal Cell Cycle by Disrupting the E2F1–DP1 Complex. J. Neurosci. 30, 5219–5228.

Zhang, H., Xue, C., Shah, R., Bermingham, K., Hinkle, C. C., Li, W., Rodrigues, A., Tabita-Martinez, J., Millar, J. S., Cuchel, M., et al. (2015). Functional Analysis and Transcriptomic Profiling of iPSC-Derived Macrophages and Their Application in Modeling Mendelian Disease. Circulation Research 117, 17–28.

Zhao, B., Wu, Q., Ye, A. Y., Guo, J., Zheng, X., Yang, X., Yan, L., Liu, Q.-R., Hyde, T. M., Wei, L., et al. (2019). Somatic LINE-1 retrotransposition in cortical neurons and non-brain tissues of Rett patients and healthy individuals. PLOS Genetics 15, e1008043.

Zheng, G. X. Y., Terry, J. M., Belgrader, P., Ryvkin, P., Bent, Z. W., Wilson, R., Ziraldo, S. B., Wheeler, T. D., McDermott, G. P., Zhu, J., et al. (2017). Massively parallel digital transcriptional profiling of single cells. Nat Commun 8, 14049.

